# PROX1 loss in adult mouse Schlemm’s canal causes permanent ocular hypertension

**DOI:** 10.1101/2025.11.13.688015

**Authors:** Sofia L. Ochoa, Hoi Lam Li, Hyeohn Kim, Zihang Yan, Natalia C. Mendonca, Pan Liu, Hyunjoo J Lee, Michael P. Vincent, Hao F. Zhang, Haiyan Gong, Evan A. Scott, Mark Johnson, Benjamin R. Thomson

**Affiliations:** Department of Biomedical Engineering, Northwestern University, Evanston, Il; Department of Ophthalmology and Department of Anatomy and Neurobiology, Boston University School of Medicine, Boston, MA; Section of Nephrology and Hypertension, Northwestern University Feinberg School of Medicine, Chicago, IL; Massachusetts Eye and Ear, Boston, Ma and Harvard Medical School, Cambridge, MA; Department of Ophthalmology, Northwestern University Feinberg School of Medicine, Chicago, IL; Department of Biomedical Engineering, NanoSTAR Institute, University of Virginia School of Medicine, Charlottsville, VA; Feinberg Cardiovascular and Renal Research Inst. Northwestern University Feinberg School of Medicine, Chicago, IL

## Abstract

Glaucoma is associated with ocular hypertension and lowering intraocular pressure is a key objective of glaucoma therapies. Recent studies have established a role for the Schlemm’s canal endothelium in this pressure increase and have shown it to have a unique, lymphatic-like, hybrid phenotype. However, the role of these lymphatic phenotypes in the adult canal remains uncertain. Long-term functional studies have been limited by systemic importance of lymphatic genes and lack of Schlemm’s canal–specific animal models. Here, we designed and validated a strategy using 4OH-tamoxifen-loaded nanocarriers to generate targeted, Schlemm’s canal specific knockout mice lacking lymphatic phenotypes. Using this system, we selectively deleted *Prox1,* the master transcription factor governing lymphatic fate. Within four weeks, intraocular pressure significantly increased, and ocular hypertension was maintained for at least 24 weeks. Unlike lymphatic vessels, which degenerate following *Prox1* deletion, Schlemm’s canal reverted to a less functional vein-like phenotype with no change in size or morphology. These results highlight the utility of nanocarriers for tissue-specific genetic recombination and demonstrate that changes in lymphatic phenotypes alter intraocular pressure, providing new targets for glaucoma therapy. Moreover, as we found that PROX1 was downregulated with age in human Schlemm’s canal, these canal-specific conditional *Prox1* knockout mice are a valuable new adult-onset model of ocular hypertension that captures key features of age-related human disease.

## Main

Glaucoma is a leading cause of vision loss worldwide, affecting an estimated 64 million people and leaving 3.6 million blind^1^. While loss of retinal ganglion cells is directly responsible for vision loss in this disease, elevated intraocular pressure (IOP) is the only modifiable risk factor for development and progression of primary open-angle glaucoma (POAG), the most prevalent form of the disease. Pharmacological reduction of IOP slows disease progression and lowers glaucoma risk in eyes with ocular hypertension^2,3^. However, even for patients on maximal medical therapy, progressive loss of visual field still occurs in some patients, indicating that current pharmacological treatment is not always sufficient and highlighting the need for new therapeutic approaches^4,5^. This is likely due to (i) the inability of current therapeutics to continuously keep IOP at levels low enough to protect the optic nerve and (ii) the lack of treatments that directly address the underlying pathology causing ocular hypertension.

The optically-clear aqueous humor flows through the anterior segment of the eye providing nutrition and removing metabolic wastes, and leaves the eye primarily through an outflow pathway comprised of the trabecular meshwork and Schlemm’s canal – a unique vessel adjacent to the iridocorneal angle that collects aqueous humor – and emptying into the collector channels and aqueous veins ^6,7^. The resistance to flow through this pathway generates IOP, and in glaucoma, this flow resistance is elevated, causing ocular hypertension. A significant fraction of the total resistance is generated as aqueous humor passes through the deeper regions of the trabecular meshwork and/or the basal lamina of Schlemm’s canal before entering the canal through pores in its endothelial inner wall. These pores modulate outflow resistance^8^ and their density is reduced in glaucoma^9,10^. It has recently been reported that this decreased pore density is negatively correlated with increased stiffness of the Schlemm’s canal endothelium in glaucomatous eyes^11,12^. Thus, it is likely that physical properties of Schlemm’s canal cells themselves, and/or their underlying basal lamina, play a central role in modulating resistance and IOP homeostasis^13^. However, how Schlemm’s canal endothelial stiffness, pore number, and outflow resistance are regulated at the molecular level, and how they become dysregulated to increase resistance in POAG remain topics of ongoing research.

Schlemm’s canal is a unique vessel with characteristics of lymphatics, such as a discontinuous basement membrane and basal to apical transendothelial flow, and those of a venule including a continuous network of cell-cell junctions^14,15^. Like other hybrid vessels^16^ in the kidney^17^, nasal mucosa^18^, placental spiral arteries^19^, and penis^20^, Schlemm’s canal is derived from blood vascular endothelium and acquires a lymphatic-like phenotype during development^21^. However, how these lymphatic characteristics impact function remains unclear. Lymphatic capillaries are highly permeable as befits their function^22^. While Schlemm’s canal endothelial cells lack the unique junctional morphology of lymphatic capillaries^21,23^, the canal inner wall has one of the highest hydraulic conductivities of any endothelium in the human body^24^, suggesting lymphatic gene expression may facilitate increased permeability.

Schlemm’s canal’s lymphatic-like “hybrid” vascular phenotype is defined by expression of the lymphatic master transcription factor PROX1^25–28^. Deletion or blockade of PROX1 or the typically lymphatic receptor tyrosine kinase VEGFR3 (also known as FLT4) during development results in an attenuated canal with disorganized morphology ^25,29–31^. However, it remains unclear whether PROX1 regulates the same transcriptional targets in the canal as it does in lymphatic vessels, and if it has a continuing role once development is complete. Other developmental pathways, including VEGFA-VEGFR2 and ANGPT1-TEK, regulate IOP homeostasis throughout life^21,32–35^ and lymphatic gene expression is maintained in the mature Schlemm’s canal, supporting continued functional importance^21,26,36^. As the elevated pressure characteristic of glaucoma is associated with a loss of tissue permeability, we hypothesized that this might be related to lymphatic character of the Schlemm’s canal endothelium and its regulation by PROX1. While a previous report did not observe IOP elevation 2 weeks after endothelial *Prox1* deletion in adult mice ^29^, these animals died within 3 weeks, and this prior study could not conclusively determine if lymphatic phenotypes were dispensable in the adult canal or if dysfunction developed over a longer timeline, consistent with human glaucoma^29^.

We have previously developed a highly specific Schlemm’s canal-targeting nanocarrier platform comprised of the amphiphilic block copolymer poly(ethylene glycol)-*b*-poly(propylene sulfide) (PEG-*b*-PPS)^37^ for drug delivery to the canal endothelium^38,39^. Here, we adapted the PEG-*b*-PPS platform to selectively deliver 4-OH-Tamoxifen (4OHT), a highly active tamoxifen metabolite that is commonly used to induce recombination in CreERT2-expressing mouse strains^40^. While 4OHT readily induces recombination, due to its low aqueous solubility it is typically delivered using hydrophobic solvents that can damage delicate ocular tissues if delivered locally^41^. We optimized PEG-*b*-PPS nanocarriers to selectively deliver 4OHT and induce tissue-specific, time-dependent Cre-mediated recombination in Schlemm’s canal endothelial cells of the mouse eye, without vehicle-induced toxicity or recombination in other tissues. In addition, expression of Cre recombinase itself is increasingly understood to cause cellular toxicity that can lead to confounding phenotypes^40^. By delivering 4OHT nanocarriers to a single eye, this system allowed us to induce deletion within an individual Schlemm’s canal and control for any Cre-induced phenotypes independent of gene deletion by injecting empty nanocarriers in contralateral eyes to generate same-animal controls.

Following validation, these targeted 4OHT-loaded nanocarriers were used to ablate Schlemm’s canal lymphatic phenotypes via deletion of *Prox1* from adult Schlemm’s canal, bypassing lethal systemic phenotypes. Within 4 weeks, we observed specific, reproducible IOP elevation that persisted throughout life, thus identifying a critical role for PROX1 and Schlemm’s canal’s lymphatic phenotype in IOP homeostasis while also generating a new adult-onset ocular hypertensive mouse model with sustained IOP elevation.

### Schlemm’s canal endothelial cells have a hybrid lymphatic-like phenotype that is mediated in both mouse and human by PROX1

The hybrid identity of Schlemm’s canal endothelial cells has been widely reported in the mouse eye at both mRNA and protein levels^21,26,28,36,42,43^. In the human eye, while lymphatic gene expression has been identified by single cell RNA sequencing ^42,44,45^, it has been less well characterized at the protein level where conflicting reports have been published^26,46^. To verify the lymphatic identity of this endothelium and determine if it is regulated by PROX1, we first performed immunostaining of mouse and human canals. Confocal microscopy of mouse eye whole mounts (Fig. 1a) and cryosections from human corneal rim tissue obtained after transplant surgery (Fig. 1b) confirmed that the canal of both species expressed lymphatic markers PROX1 and FLT4, but not LYVE1 (Supplementary Fig. 1) confirming their hybrid identity. PROX1 expression was negatively correlated with donor age (Supplementary Fig. 2a), consistent with findings in the mouse eye^47^.

**Figure 1.**
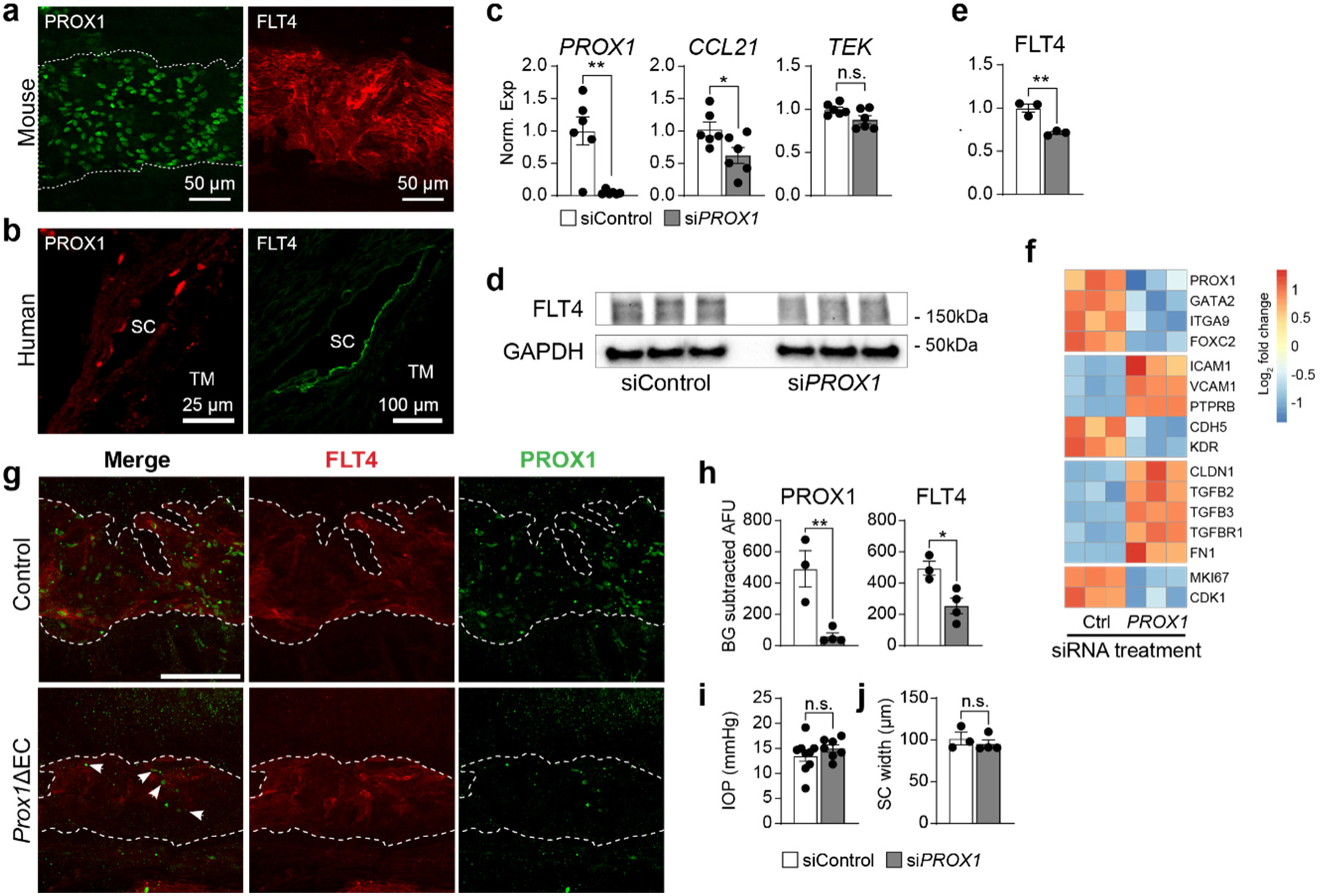
**Human and mouse SC endothelial cells share a lymphatic-like hybrid phenotype mediated by PROX1. a-**Whole mount immunostaining and cryo cross-sections of mouse (**a**) and human (**b**) Schlemm’s canals revealed robust expression of the lymphatic markers PROX1 and FLT4. (c) *PROX1* and *CCL21* expression were reduced in primary human SC endothelial cells treated with PROX1 siRNA, while levels of the universally expressed endothelial gene *TEK* were unchanged when measured by real-time qPCR (siControl n = 6, si*PROX1* n = 6). (**d**, quantified in **e**) Western blot revealed reduced expression of FLT4 protein in siPROX1-treated human SC cells (n = 3 per group). (**f**) Reduced expression of lymphatic genes and increased expression of blood endothelial genes were detected in si*PROX1*-treated SC cells by RNA sequencing, accompanied by increased expression of TGFB-signaling genes and reduction in cell proliferation markers (n = 3 per group). (**g**, quantified in **h**) Confocal microscopy of eye flat mounts revealed reduced PROX1 and FLT4 expression in *Prox1*^flox/flox^; *Cdh5*^CreERT2^ mice 4 weeks after tamoxifen induction at 8 weeks of age (*Prox1*ΔEC). While *Prox1* deletion was generally robust, some mosaicism was observed, and a small number of PROX1-positive nuclei were observed in *Prox1*ΔEC SC (white arrowheads, *Prox1*ΔEC n = 3, Control n = 4). Dashed lines in **g** outline Schlemm’s canal. (**i**) No change in IOP (*Prox1*ΔEC n = 9, Control n = 7) or (**j**) Schlemm’s canal width (*Prox1*ΔEC n = 3, Control n = 4) was measured 4 weeks after tamoxifen induction. * p <0.05, ** p <0.01 as determined by 2-tailed unpaired Student’s t test. Error bars in **c**, **e**, **h** and **i** indicate ±SEM, while each point denotes an independent biological replicate.

As PROX1 directly regulates the fate of lymphatic endothelial cells, we speculated that it played a similar role in Schlemm’s canal. To test this directly, we treated primary human Schlemm’s canal cells with siRNA targeting *PROX1*. 72 h after treatment, real-time rtPCR was used to confirm *PROX1* knockdown (Fig. 1c). Expression of *CCL21* and *FLT4*, additional lymphatic markers expressed by Schlemm’s canal endothelial cells (Fig. 1c-e)^26^, were also suppressed, suggesting loss of PROX1 resulted in a general reduction of lymphatic gene expression.

Expression of *TEK*, an endothelial receptor tyrosine kinase expressed in both blood and lymphatic endothelial cells, was unchanged. RNA sequencing revealed downregulation of additional canal-expressed lymphatic genes in si*PROX1*-treated cells, including *ITGA9* and *GATA2* (Fig. 1f, Supplemental dataset 1). In contrast, a subset of blood-endothelial-specific genes including *ICAM1*, *VCAM1* and *PTPRB* were upregulated. Together this data suggested that, as in lymphatic endothelial cells ^48^, the lymphatic transcriptional program in Schlemm’s canal endothelial cells was mediated by PROX1.

To understand the role of PROX1 in mediating the Schlemm’s canal hybrid phenotype *in vivo*, we next turned to an endothelial-specific *Prox1* knockout mouse model^49^. *Prox1*^flox/flox^; *Cdh5-*CreERT2 mice were generated and induced with tamoxifen at 8 weeks of age to obtain animals lacking *Prox1* in all endothelial cells, including those in Schlemm’s canal (*Prox1*ΔEC). Four weeks after induction, immunostaining revealed robust PROX1 ablation, although some mosaicism was observed and occasional endothelial cells retained PROX1 (Fig. 1g, quantified in h). FLT4 staining was also reduced, consistent with our *in vitro* data and the well-known role of PROX1 in regulating FLT4 expression in lymphatic endothelial cells^48^. IOP measurement by rebound tonometry (Fig. 1i), revealed normal IOP, and no change in Schlemm’s canal size was observed when measured using immunofluorescence confocal microscopy in canal whole mounts (Fig. 1j).

Lack of IOP elevation two weeks after whole endothelial *Prox1* deletion has been previously reported^29^ and, as in those reports, we were unable to track the IOP of *Prox1*ΔEC mice beyond 4 weeks due to intestinal hemorrhage and lethality, likely caused by the role of lymphatic vessels in maintaining the gut epithelium^47,50^. While these data confirmed PROX1 regulated Schlemm’s canal’s lymphatic gene expression, we remained skeptical that the hybrid identity was unnecessary for IOP homeostasis—especially as the finding that PROX1 expression decreased with donor age is consistent with an age-related increase in outflow resistance that occurs in humans^51^. Thus, we speculated that the short survival time of *Prox1*ΔEC mice was simply insufficient to observe an IOP phenotype, or that whole-endothelial *Prox1* deletion caused confounding phenotypes that lowered IOP or prevented accurate tonometric measurements.

To bypass systemic phenotypes, we determined that Schlemm’s canal-specific deletion would be required. However, no canal-specific Cre-expressing mouse line has been described, necessitating the development of a new approach. We therefore employed the PEG-*b*-PPS nanocarrier delivery system previously developed by our group in order to selectively deliver the active metabolite 4OHT directly to Schlemm’s canal endothelial cells of *Prox1*^flox/flox^; *Cdh5CreERT2* mice to induce Schlemm’s canal-specific cre-mediated recombination.

### Targeted 4OH-tamoxifen nanocarriers specifically induce gene deletion in Schlemm’s canal endothelial cells

Modifying a PEG-*b*-PPS delivery system that was previously developed by our group^37–39^, we designed novel targeted nanocarriers to selectively deliver 4OHT to Schlemm’s canal endothelial cells *in vivo* to achieve SC-specific, Cre-mediated gene deletion. Micellar nanocarriers were generated, loaded with 4OHT, and decorated with VEGFC-derived peptide-lipid targeting constructs (PG48), which we have previously optimized for FLT4 binding and selective delivery to the Schlemm’s canal endothelium (Fig. 2a)^39,52^. Unlike unmetabolized tamoxifen that must circulate for processing by cytochrome P450 in the liver^53^, 4OHT is active immediately in cells where it is delivered, making it the optimal cargo for this application. 4OHT nanocarriers exhibited a stable monodisperse particle size distribution (PDI ≤ 0.2) with a diameter of 17-22 nm when assessed by cyro-electron microscopy and dynamic light scattering (DLS, Fig. 2b, Supplementary Fig. 3a and Table 1). Zeta-potential was determined by electrophoretic light scattering (ELS), indicating a neutral surface charge (Table 1). Spherical morphology was confirmed by cryo-transmission electron microscopy (Cryo-TEM) and small angle x-ray scattering (SAXS, Fig. 2 b,c) and was unaffected by addition of 4OHT and targeting peptides.

**Figure 2.**
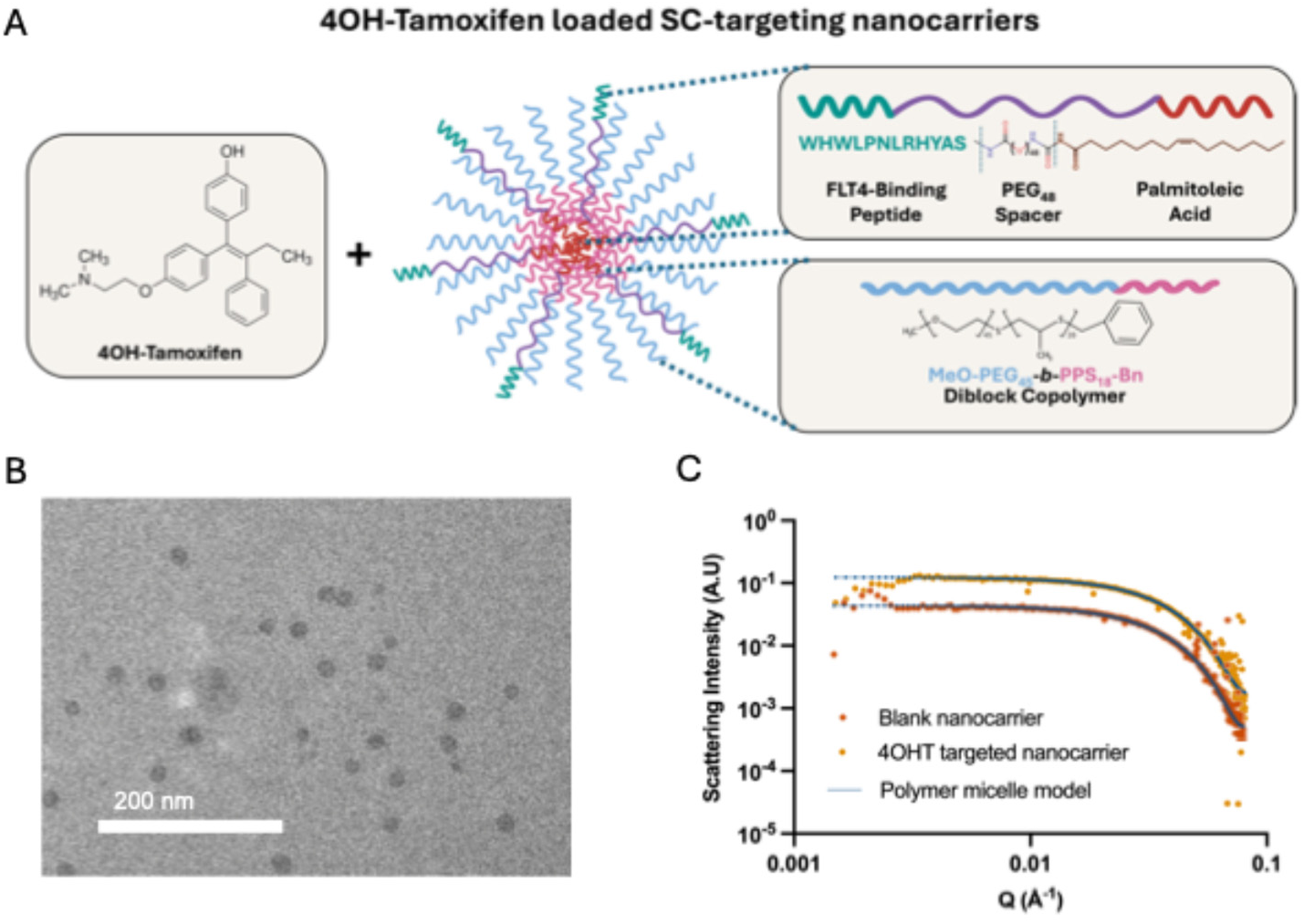
Material characterization of targeted and untargeted 4OH-Tamoxifen loaded nanocarriers (Targeted 4OHT/Untargeted 4OHT) (A) Schematic illustrating how PEG-*b*-PPS targeted nanocarriers are generated, loaded with 4-OH-Tamoxifen. (B) Micelle morphology for 4-OHT nanocarriers was characterized by (B) cryogenic transmission electron microscopy (Cryo-TEM) and (C) small angle X-ray scattering (SAXS, n= 2).

**Table 1.**
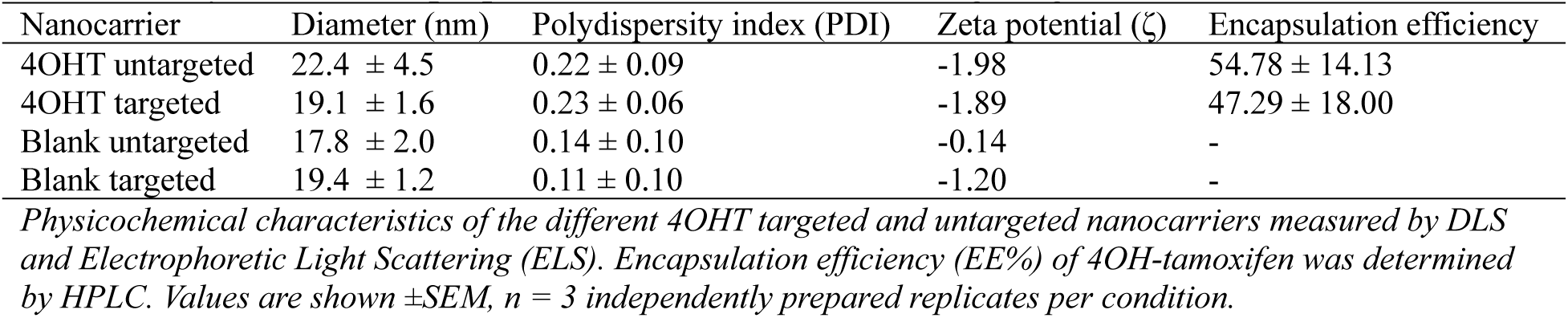
Physiochemical properties of Schlemm’s canal targeting nanocarriers.

|  |  |  |  |  |
| --- | --- | --- | --- | --- |
| Nanocarrier | Diameter (nm) | Polydispersity index (PDI) | Zeta potential ($\zeta$) | Encapsulation efficiency |
| 4OHT untargeted | 22.4 ± 4.5 | 0.22 ± 0.09 | -1.98 | 54.78 ± 14.13 |
| 4OHT targeted | 19.1 ± 1.6 | 0.23 ± 0.06 | -1.89 | 47.29 ± 18.00 |
| Blank untargeted | 17.8 ± 2.0 | 0.14 ± 0.10 | -0.14 | - |
| Blank targeted | 19.4 ± 1.2 | 0.11 ± 0.10 | -1.20 | - |

*Physicochemical characteristics of the different 4OHT targeted and untargeted nanocarriers measured by DLS and Electrophoretic Light Scattering (ELS). Encapsulation efficiency (EE%) of 4OH-tamoxifen was determined by HPLC. Values are shown ± SEM, n = 3 independently prepared replicates per condition.*

Encapsulation efficiency of 4OHT was 47-54% (Table 1) with a resulting concentration of 80-120 µg/mL encapsulated 4OHT in injectable formulations. No cytotoxicity was observed in cultured HUVEC cells treated with nanocarrier concentrations up to 2μM 4OHT (Supplementary Fig. 3b). Validating the targeting approach, flow cytometry demonstrated that VEGFC-derived SC targeting peptides increased uptake of DiI-labeled nanocarriers by primary human Schlemm’s canal cells but not in HUVECs, which do not express FLT4 (Supplementary Fig. 3c). Although tamoxifen has been delivered via nanoparticles to assess in vivo nanoparticle targeting efficiency^54^ and as a chemotherapeutic for cancer treatment^55^, to the best of our knowledge, this is the first application of 4OHT-loaded nanocarriers for cell selective modulation of gene expression to model a disease state.

### 4OH-tamoxifen nanocarriers specifically induce gene deletion in Schlemm’s canal *in vivo*

To determine if the newly designed FLT4-targeted 4OHT nanocarriers could induce high efficiency, specific recombination in Schlemm’s canal, we generated cohorts of *Rosa26*^mTmG^; *Cdh5*-CreERT2 reporter mice and administered nanocarriers via intracameral injection (Fig. 3a). Nanocarriers decorated with FLT4-specific Schlemm’s canal-targeting peptide were delivered to one eye of each animal and untargeted nanocarriers in the other. Seven days later, while we observed recombined endothelial cells in Schlemm’s canals receiving either targeted or untargeted nanocarriers, efficacy of targeted nanocarriers was markedly higher, confirming the success of our targeting approach (Fig. 3b).

**Figure 3.**
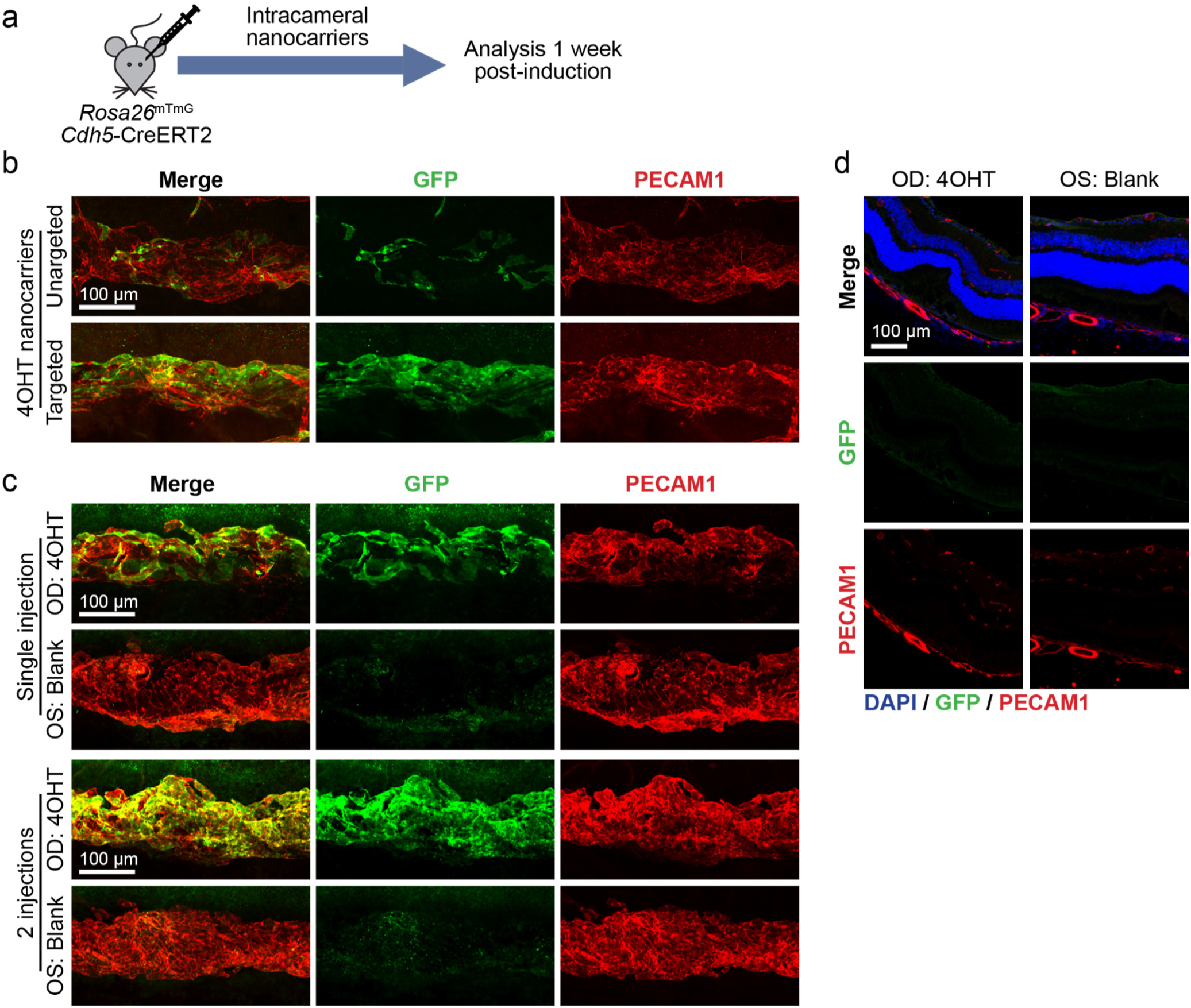
4OH tamoxifen loaded, SC targeted nanocarriers induce robust recombination in SC endothelial cells. (**a**) Schematic of experimental timeline used for nanocarrier induction. (**b**) 7 days after intracameral infusion of 4OHT-loaded nanocarriers, robust recombination was observed in eyes treated with nanocarriers labeled with FLT4 targeting peptide, while only sporadic recombination was observed in eyes receiving untargeted 4OHT nanocarriers. (**c**) SC-specific recombination efficiency was further increased by a second injection of FLT4-targeting nanocarriers 24h after the first. (**d**) confirming specificity of the system, no recombination was observed in retinal or choroidal capillaries of eye cryosections prepared from *Rosa26*^mTmG^; *Cdh5*-CreERT2 mice 7 days after receiving two intracameral injections of targeted 4OHT or blank nanocarriers.

We next tested repeat nanocarrier injections as a tool to increase recombination efficiency (Fig. 3c). Following a second injection 24 h after the first, we observed nearly complete labeling of the Schlemm’s canal endothelium. Despite robust recombination within the eye receiving 4OHT nanocarriers, no recombination was observed in Schlemm’s canal of contralateral eyes that received empty control nanocarriers, or within retinal or choroidal capillaries of either eye (Fig. 3d), confirming specificity of the targeting approach. Mosaic recombination was seen in distal outflow vessels, suggesting some nanocarriers flowed through the canal without uptake by Schlemm’s canal endothelium, and in limbal lymphatic capillaries, consistent with their known expression of FLT4 (Supplementary Fig. 4a,b). Outside of the eye, while we observed GFP expression in some endothelial cells of the liver, kidney and lung (Supplementary Fig. 5), recombination levels were similar to those in untreated animals, indicating that recombination in these tissues was due to leakiness of Cre activity and not off target nanocarrier delivery.

### Schlemm’s canal-specific Prox1 deletion caused ocular hypertension in adult mice

Following validation of targeted 4OHT nanocarriers, we returned to our original question—the long-term role of PROX1 in maintaining IOP homeostasis and lymphatic identity of Schlemm’s canal. We generated cohorts of *Prox1*^flox/flox^; *Cdh5*-CreERT2 mice with Cre-negative littermate controls. At 8 weeks of age, Schlemm’s canal-specific *Prox1* deletion was induced in randomized eyes with Schlemm’s canal targeted 4OHT nanocarriers. Contralateral eyes received identical injections of empty (blank) nanocarriers as a control (Fig. 4a). At baseline, we observed no difference in IOP between eyes (ΔIOP_baseline_ = 0.2±0.60 mmHg, p >0.9), but beginning 4 weeks after induction, IOP in 4OHT-treated eyes was significantly elevated in comparison with same-animal control eyes (Fig. 4b, ΔIOP_4wks_ = 4.5±0.8 mmHg, p < 0.005). No IOP difference was observed between eyes in Cre-littermates insensitive to tamoxifen (ΔIOP_4wks_ =-0.6±0.5 mmHg, p >0.9). IOP remained elevated for 18 weeks, as long as the animals were maintained. In an independent replication cohort, elevated IOP was observed for 24 weeks, again as long as the animals were maintained (Supplementary Fig. 6, ΔIOP_24wks_ = 4.4±1.3 mmHg, p < 0.001).

**Figure 4.**
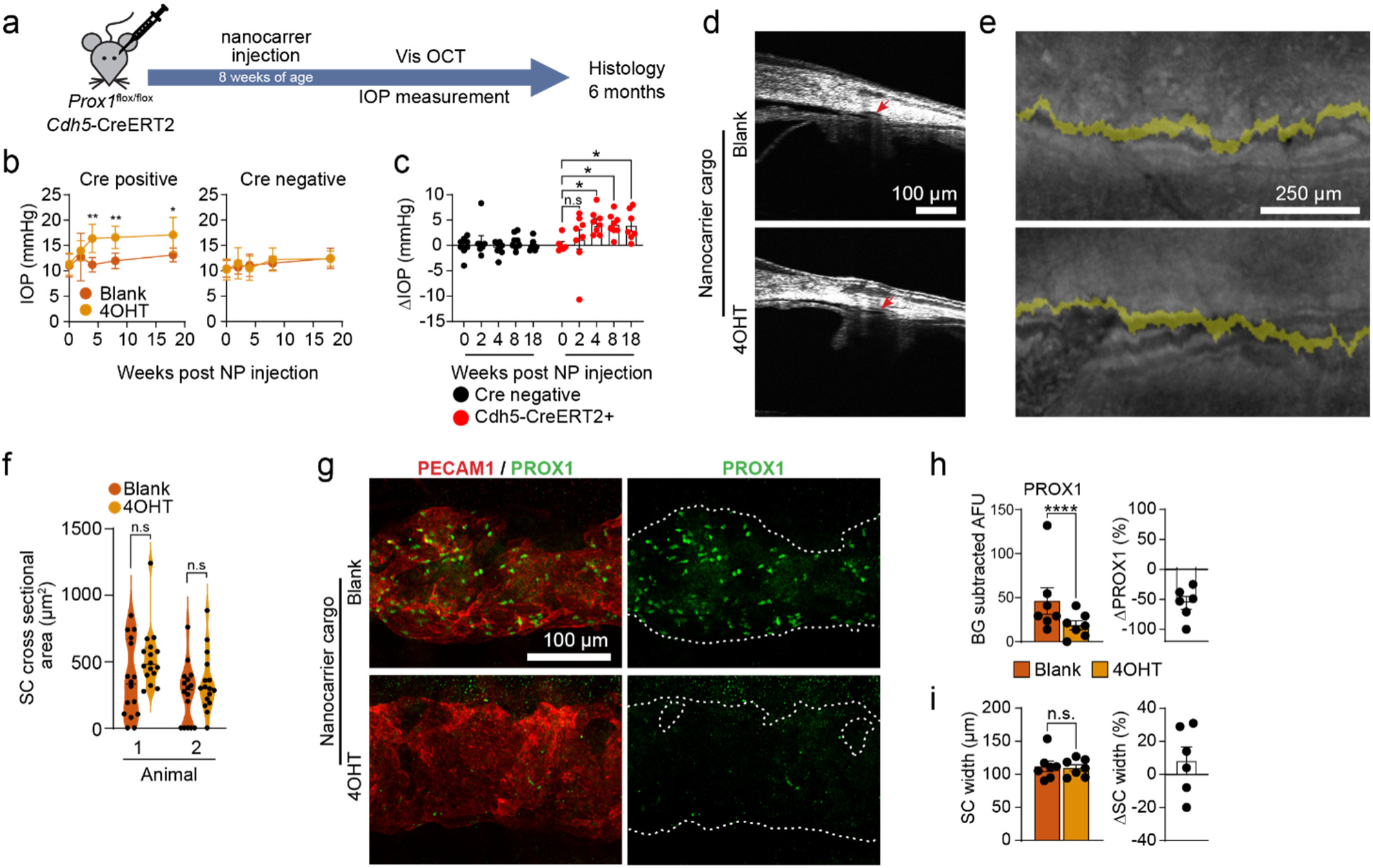
**SC-specific *Prox1* deletion leads to ocular hypertension**. (**a**) Schematic of experimental timeline used for nanocarrier induction of *Prox1* deletion. (**b**, **c**) 4 weeks after nanocarrier induction, IOP elevation was observed in eyes of *Prox1*^flox/flox^;*Cdh5*-CreERT2 mice receiving 4OH tamoxifen (4OHT) nanocarriers in comparison to contralateral eyes receiving empty (blank) nanocarriers. IOP elevation persisted throughout the duration of the experiment. No IOP elevation was seen in Cre-negative mice treated with 4OHT nanocarriers. (Cre+, n = 8, Cre-, n = 8). (**d**) In vivo visible light-OCT imaging 12 weeks after nanocarrier-mediated *Prox1* deletion revealed no change in canal size on B-scans, or (**e**) longitudinal reconstructions (pseudo-colored in yellow) of Schlemm’s canal. (**f**) Comparison of luminal area by 16 individual OCT B scans captured around the circumference of Schlemm’s canal showed no difference in canal size between matched 4OHT and blank nanocarrier-treated eyes of *Cdh5*-CreERT2-positive mice. (**g**) PROX1 expression and canal morphology were examined in Schlemm’s canal flat mounts collected 6 months after nanocarrier induction by confocal microscopy. Dashed lines in PROX1 panels indicate the outline of PECAM1-positive Schlemm’s canal. (**h**) Compared with contralateral control eyes, PROX1 expression was significantly reduced in eyes treated with 4OHT nanocarriers. (**i**) No change in Schlemm’s canal size was observed in 4OHT nanocarrier-treated eyes (n = 6). * p < 0.05, ** p < 0.01, **** p < 0.0001 as determined by 2-way ANOVA followed by Bonferroni posttests (**b**, **c** and **f**) or 2-tailed paired Student’s t test (**h** & **i**). Error bars in **b**, **c**, **h**, and **i** indicate ±SEM, while each point represents an independent biological replicate. Each point in **f** represents a single measurement captured along the length of Schlemm’s canal, while each violin represents a single eye.

Surprisingly, given increased IOP and the central role of PROX1 in Schlemm’s canal development, *in-vivo* visible light optical coherence tomography (vis-OCT) imaging performed 12 weeks after induction revealed no difference in Schlemm’s canal size or morphology between PROX1 knockout and same animal contralateral control eyes (Fig. 4d-e, quantified in f).

Following longitudinal IOP measurements, enucleated eyes were stained and prepared for immunofluorescent imaging (whole mounts) and for light and electron microscopy (cross-sections). Compared to contralateral controls, confocal microscopy confirmed reduced PROX1 immunostaining in canals of 4OHT nanocarrier-treated eyes (56±10.8% reduction, p <0.001; Fig. 4g, quantified in h). Schlemm’s canal size, as measured by PECAM1-positive immunofluorescence, was unchanged (Fig. 4i). At 12 weeks of age, light microscopy in a separate group of animals revealed no differences in gross canal morphology or geometric parameters between *Prox1* knockout and contralateral control eyes (Fig. 5a-d). Similarly, when imaged using transmission electron microscopy (Fig. 5e), we did not observe differences in the morphology of Schlemm’s canal inner wall or juxtacanalicular connective tissue. Giant vacuoles were observed in the Schlemm’s canal of both *Prox1* knockout and control eyes.

**Figure 5.**
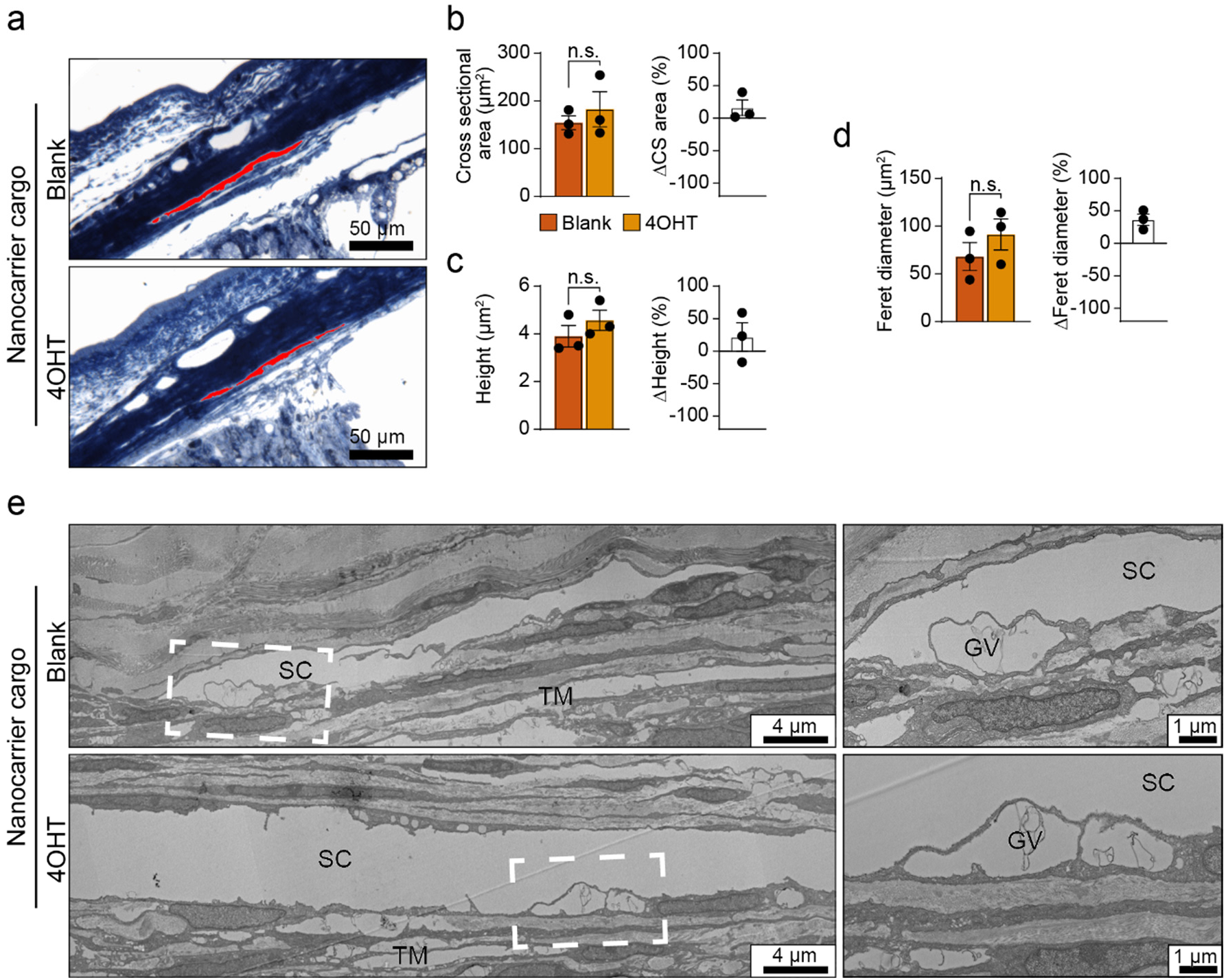
Light and electron microscopy revealed normal Schlemm’s canal morphology in *Prox1* knockout SC. (**a**) Representative toluidine blue-stained semithin cross-sections of Schlemm’s canal from *Prox1*^flox/flox^; *Cdh5*-CreERT2 eyes 12 weeks after treatment with blank or 4OHT nanocarriers. Semithin sections were imaged by light microscopy and used to measure (**b**) SC cross sectional area, (**c**) canal height (depth) and (**d**) Feret diameter (longest axis; n = 3 pairs of eyes from *Cdh5*-CreERT2-positive nanocarrier-treated animals). Schlemm’s canal lumen indicated by red shading. When compared to same-animal contralateral control eyes, no significant difference in canal size or morphology was observed. (**e**) Representative TEM images from the inferior quadrant of matching eyes treated with 4OHT or blank nanocarriers from a *Prox1*^flox/flox^; *Cdh5*-CreERT2 mouse revealed no differences in SC inner wall or juxtacanalicular meshwork morphology 12 weeks after nanocarrier injection. Giant vacuoles (GV) were observed in the inner wall of both 4OHT-treated and contralateral control eyes. Statistical comparisons in **b**-**d** were performed using a Bonferroni-corrected 2-tailed, paired Student’s t test. Error bars in **b**, **c**, and **d** indicate ±SEM, while each point represents an independent biological replicate.

These findings suggested that increased IOP seen following *Prox1* deletion was due to alterations in outflow function of the targeted Schlemm’s canal cells and/or in their communication and interactions with neighboring cells or extracellular matrix, rather than canal degeneration or gross morphological changes. Furthermore, these findings confirmed the efficacy of FLT4-targeted 4OHT nanocarriers for gene deletion within Schlemm’s canal and demonstrated that PROX1-mediated lymphatic hybrid phenotypes were essential for IOP homeostasis.

In contrast to the long-term persistence of Schlemm’s canal following *Prox1* deletion, we observed loss of limbal lymphatic capillaries in 4OHT nanocarrier-treated eyes, which are targeted by our FLT4-based strategy in addition to Schlemm’s canal (Supplementary Fig. 7), consistent with their requirement for ongoing PROX1 activity ^48,56^. Normal PROX1 expression was seen in a small number of remaining lymphatics suggesting that deletion efficacy in these vessels was lower than in Schlemm’s canal—perhaps because the limbal lymphatics do not connect to the anterior chamber and lacked a direct route for nanocarrier entry.

### Endothelial-specific Flt4 knockout mice induced as adults have normal IOP

While PROX1 has many direct transcriptional targets, one of the best characterized is *Flt4*. *PROX1* knockdown in cultured lymphatic endothelial cells^48^ and in primary human Schlemm’s canal cells (Fig. 1) led to reduction of *FLT4*, and *PROX1* overexpression drives ectopic *FLT4* expression in blood vascular endothelial cells^48^. In mouse lymphatic vessels, *Prox1* deletion leads to loss of FLT4 and the lymphatic phenotype overall^57,58^. Accordingly, as FLT4 is essential for Schlemm’s canal development and its ligand VEGFC induces angiogenic sprouting from the adult canal^26^, we speculated that FLT4 downregulation could be the cause of IOP elevation in *Prox1* knockout eyes. Therefore, we generated endothelial-specific *Flt4* knockout mice using *Cdh5*-CreERT2 (*Flt4*ΔEC), and induced deletion by tamoxifen injection at 8 weeks of age (Fig. 6a).

**Figure 6.**
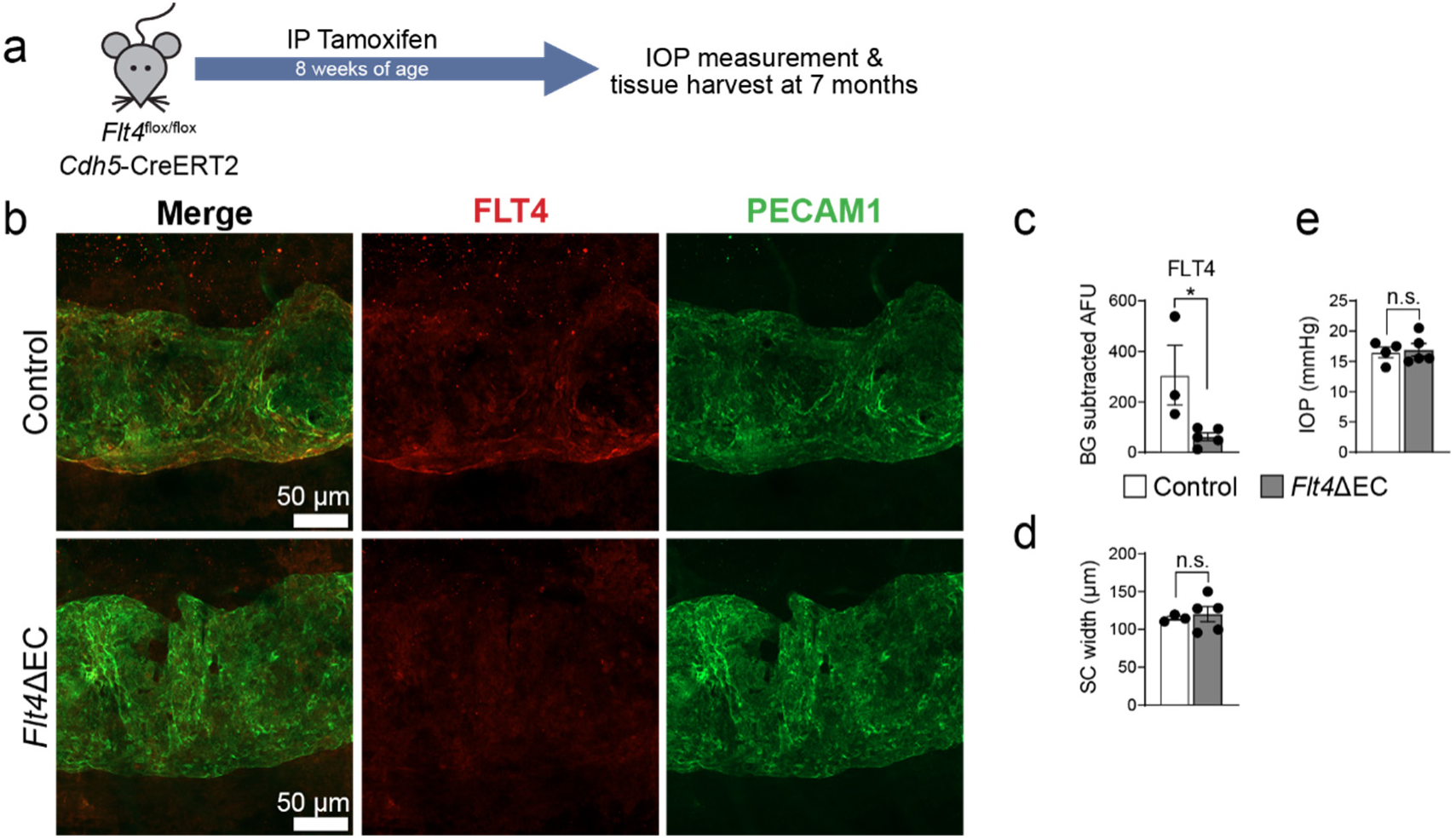
FLT4 is not required for SC maintenance or IOP homeostasis in adult mice. (**a**) Experimental outline used for generation and analysis of endothelial-specific *Flt4* knockout mice (*Flt4*ΔEC). (**b**, quantified in **c** & **d**) Confocal microscopy revealed loss of FLT4 immunostaining and similar PECAM1-positive SC size 5 months after *Flt4* deletion. Control, n = 5, *Flt4*ΔEC, n = 3. (**e**) No difference in IOP was observed in between *Flt4*ΔEC mice and control littermates at 7 months of age. Control, n = 5, *Flt4*ΔEC, n = 4. Statistical comparisons in **c**-**e** were performed using a 2-tailed Student’s t test. * p < 0.05. Error bars indicate ±SEM, while each point represents an independent biological replicate.

Twenty weeks after tamoxifen administration, confocal microscopy revealed loss of FLT4 immunostaining from Schlemm’s canal in *Flt4*ΔEC mice (Fig. 6b, quantified in Fig. 6c) with no change in PECAM1-positive Schlemm’s canal size or morphology (Fig. 6d). Surprisingly, no difference in IOP was observed between *Flt4*ΔEC mice and Cre negative control littermates (ΔIOP = 0.4 mmHg, p > 0.7; Fig. 6e). These data indicated that, while FLT4 is essential for Schlemm’s canal development^26^, it was not responsible for the IOP elevation in *Prox1*ΔSC eyes and was not required for IOP homeostasis in adult mice.

## METHODS

### Study Approval

Animal experiments were approved by the Animal Care and Use Committee at Northwestern University (Evanston, IL) under animal protocol numbers IS00020857 and IS00022641 and comply with ARVO guidelines for care and use of vertebrate research subjects in Ophthalmology research.

### Animal generation and husbandry

During all experiments, mice were housed at the Center for Comparative Medicine of Northwestern University (Chicago, IL USA). Animals were provided with unlimited access to water and standard rodent diet (Teklad #7912) and maintained on a standard 12-hour lighting cycle at a temperature of 21-23° and relative humidity of 30-70%. To generate litters of *Rosa26*^mTmG^; *Cdh5*-CreERT2 mice for analysis, gt(ROSA)26Sortm4(ACTB-tdTomato,-EGFP)Luo/J (*Rosa26*^mTmG^, Jax strain #007576)^59^ mice were crossed with Tg(Cdh5-cre/ERT2)^1Rha^ animals carrying the *Cdh5*-CreERT2 transgene ^60^. *Prox1*^flox/flox^ mice were a generous gift of Dr. Guillermo Oliver (Northwestern University, Chicago, IL) ^49,61^, and *Flt4*-floxed mice were a gift of Dr. Susan Quaggin (Northwestern University, Chicago, IL) ^62^.

Throughout the study, animals were maintained on a mixed genetic background, and animals of both sexes were included in all experiments. Mice were genotyped by PCR using previously published primers.

### Primary human Schlemm’s canal cell culture

Primary human Schlemm’s canal cells were a generous gift of Dr. W. Daniel Stamer (Duke University, Durham, NC). For siRNA experiments, normal SC cells from clones SC-68 and SC-91 were seeded in 6 well plates before transfection in triplicate groups using *PROX1*-specific siRNA (Dharmacon, Lafayette, CO, USA. #016913-00-0005) or matching scrambled siRNA control (Dharmacon #016913-00-0005) in combination with RNAimax reagent (Invitrogen, Thermo Fisher Scientific, 13778075) according to the manufacturer’s instructions. 72 hours after treatment, total RNA was collected using TRIzol reagent (ThermoFisher Scientific #15596026) and purified (RNEasy MinElute cleanup kit; Quagen, Hilden Germany) before generation of cDNA for real-time PCR or RNA sequencing.

### HUVEC cell culture

Human Umbilical Vein Endothelial Cells (HUVEC-Umbil Vein, Pooled,EGM-2,amp) from pooled donors were purchased from Lonza. All primary cells used in these studies were used at passage 4. HUVECs were cultured in endothelial cell growth basal medium-2, (EBM™-2, Lonza) supplemented with FBS and EGM™-2 Endothelial Cell Growth Medium-2 BulletKit™ (Lonza) optimized for HUVEC culture. All cells were cultured at 37°C, 5% CO_2_ in T25 flasks.

### Analysis of human corneal rims

Surplus corneal rim tissue was obtained following corneal transplant surgery, fixed (4% formaldehyde) and cryosectioned using standard techniques. Tissues recovered and preserved for transplant within 15 hours of death were accepted for analysis; tissue with longer recovery times showed no PROX1 expression (data not shown). No restrictions were placed on duration of storage in cornea culture media prior to transplant surgery. Cryosections were washed and blocked (5% donkey serum, 2.5% BSA in TBS containing 0.5% Triton X100) prior to overnight incubation with primary antibodies at 4°. Sections were then washed and incubated with appropriate alexafluor-labeled secondary antibodies. To quantify PROX1 expression within Schlemm’s canal, sections were co-stained using antibodies against PROX1 and the endothelial transcription factor ERG and QuPath software was used to quantify PROX1 fluorescence intensity in ERG-positive SC nuclei and background intensity in non-endothelial ERG-negative, DAPI-positive nuclei in other cells of the iridocorneal angle. Identically prepared sections from human pterygium tissue were used as a positive control for LYVE1 staining. Primary antibodies used: Mouse anti PROX1 (Developmental Studies Hybridoma Bank, Iowa City, IA USA, 1A6), Goat anti LYVE1 (R&D systems AF2125), Goat anti FLT4 (R&D systems, AF349), Rabbit anti ERG (Abcam, Waltham MA, USA, ab92513).

### Real-time quantitative PCR

Following RNA purification, cDNA was prepared using the iScript Kit (Bio-Rad Laboratories, Hercules CA) according to the manufacturer’s instructions. Real-time PCR was then performed using a QuanStudio3 instrument (ThermoFisher Scientific) and Power SYBR green master mix (ThermoFisher Scientific). The following primers were used: GAPDH-Fwd: 5’-AAGGTCATCCCAGAGCTGAA-3’, GAPDH-Rev: 5’-CTGCTTCACCACCTTCTTGA-3’, PROX1-Fwd: 5’-GAGCCTCCGTGGAACTCA-3’, PROX1-Rev: 5’-TGGGCACAGCTCAAGAATC-3’, TEK-Fwd: 5’-CCCCTATGGGTGTTCCTGT-3’, TEK-Rev: 5’-GCTTACAATCTGGCCCGTAA-3’, CCL21-Fwd: CGCAGCTACCGGAAGCAG-3’, CCL21-Rev: 5’-CTGCCTGAGAGCGCTTGC-3’.

### Western blot

Following siRNA transfection as described above, triplicate samples of primary human SC cells (clone SC-68) were lysed using Laemmli sample buffer containing 100 mM DTT and separated using a 4-15% acrylamide minigel (TGX, Bio-Rad Laboratories). Proteins were then transferred onto a PVDF membrane (Bio-Rad), blocked (5% bovine serum albumen in TBS pH 7.5 with 0.05% Tween 20, 1 h at room temperature) before incubation overnight with appropriate primary antibodies. Membranes were then washed (TBS containing 0.05% tween 20), incubated with appropriate HRP-conjugated secondary antibodies and visualized using ECL reagent. Images were captured using an iBright CL1500 imaging system (ThermoFisher Scientific) before densitometry was performed using ImageJ software. Primary antibodies used: Goat anti-human FLT4 (R&D Systems, Minneapolis MN, #AF349 1:100 dilution), rabbit anti human GAPDH (Cell Signalling, Danvers MA, #2118 1:10,000 dilution).

### RNA sequencing

Primary human SC cell clone SC-68 was used for RNA sequencing. Following RNA purification as described above, total RNA isolated from triplicate samples of siControl and si*PROX1*-treated cells were provided to the NUSeq core of Northwestern University Feinberg School of Medicine for library preparation (TruSeq Kit; Illumina, San Diego, CA) and sequencing on an Illumina HiSeq 4000 instrument. Reads were then aligned to the human genome and differential expression analysis was performed using DeSeq2. Differentially expressed genes were then filtered using the Benjamini-Hochberg method using a false discovery threshold of 0.05 and representative genes were selected for the heat map shown in figure 1.

### Solid-phase peptide synthesis

Standard Fmoc solid-phase peptide synthesis was performed to synthetize the FLT4-binding targeting peptide. Fmoc-N-amido-dPEG_24_ (Quanta Biodesign) were purchased for the use in synthesis of the PG48 peptide constructs. The chemical structure of the peptide is presented in figure 2 a.

### Nanocarrier generation and optimization

All polymers used for the formulation of nanocarriers were synthesized based on previously established procedures^63^. Briefly, 40 mg of polymer, 136 μg 4OH-Tamoxifen (4OHT, H6278, Millipore Sigma, St. Louis MO) and 5 μL of DiI dye (42364-100MG, Sigma-Aldrich)) were dissolved in 500 μL of tetrahydrofuran (THF). This mixture was then added dropwise to 1 mL of Phosphate Buffer Saline (PBS, pH 7.5) under constant rotation. Nanocarrier solution was left in a desiccator to remove THF overnight. FLT4 binding peptide (5% molar ratio) was dissolved in DMSO and incorporated to the nanocarriers by gentle rotation. Blank nanocarriers were mixed with an equivalent amount of DMSO without FLT4-binding peptide. Finally, the nanocarriers were purified and concentrated to a final volume of 250 mL. Drug concentration and particle size were characterized by High Performance Liquid Chromatography (HPLC) and Dynamic Light Scattering (DLS). Blank nanocarriers were synthesized by the same procedure in the absence of 4OHT.

### Quantification of 4OHT loading into nanocarriers

50 mL aliquots of nanocarrier solution were frozen and freeze-dried. Resulting powder was dissolved in methanol and precipitated at-20°C. Solution was then centrifuged (4000*g, 5 m) to separate the precipitated polymer, and supernatant was collected and analyzed. 4OHT concentration was determined using HPLC, with calibration against a 4OHT concentration series (Supplementary Fig. 3D) prepared in methanol. Absorbance was measured at 235 nm. Data was obtained from three replicate samples, and HPLC was conducted using a C18 XDB-Eclipse column (Agilent) with a static mobile phase of acetonitrile (ACN) and 0.1% TFA HPLC water (85:15). 4OHT encapsulation efficiency (EE%) provided as a percentage was calculated as 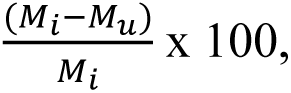 where M_i_ in the input mass of 4OHT and M_u_ is the mass of unencapsulated 4OHT.

### Quantification of nanocarrier size, polydispersity, and zeta potential

To determine particle diameter size and polydispersity index (PDI), 10 mL aliquots of targeted and untargeted nanocarriers were diluted in PBS at a 10% v/v concentration and characterized by DLS using a Zetasizer Nano instrument (Malvern Instruments) equipped with a 4 mW He-Ne 633 nm laser. Diameter of spherical nanocarriers and PDI were determined based on the intensity average size distributions from three independently prepared samples (n=3). Zeta ζ potential was measured using a Zetasizer Nano device (Malvern Instruments). Average ζ potential for each nanocarrier was calculated based on measurements from three separate formulations.

### Cryogenic Transmission Electron Microscopy

Copper grids (200 mesh) with a lacey carbon membrane (Cat# LC200-Cu-100; Electron Microscopy Sciences, Hatfield, PA) were glow-discharged in a Pelco easiGlow Discharge Cleaning System. A sample volume of 4 mL was added to the grid and blotted for 5 seconds with a blot offset of 0.5 mm and plunged into liquid ethane within a FEI Vitrobot Mark III plunge freezing instrument. Samples were imaged using a JEOL JEM1230 Lab5 emission TEM (JEOL USA) operating at 100 kEV.

### Small-Angle X-ray Scattering (SAXS)

Micelle morphology was confirmed and characterized by small angle x-ray scattering (SAXS) using synchrotron radiation at the DuPont-Northwestern-Dow Collaborative Access Team (DND-CAT) beam-line at the Advanced Photon Source at the Argonne National Laboratory. A core-shell sphere model was fit to the scattering profile of blank nanocarriers (without peptide) as well as 4OH-Tamoxifen nanocarriers displaying PG48 targeting peptide construct at 5% molar ratio (peptide/polymer).

### In vitro biocompatibility of 4OHT nanocarriers

The 3-(4,5-Dimethylthiazol-2-yl)-2,5-Diphenyltetrazolium Bromide (MTT) assay was employed to measure cell viability following incubation with nanocarriers. HUVECs, at a concentration of 5×10^5^ cells/mL determined by cell counting, were seeded into U-bottom 96-well plates, with 200 µL of cell suspension per well. Each well received nanocarriers in 3 replicates, prepared in PBS with a final 4OHT concentrations of 0, 0.5, 1, and 2 μM. Cells were incubated with nanocarriers for 24 hours at 37 °C and 5% CO_2_. MTT reagent, prepared at 5 mg/mL in PBS, was added to each well (20 µL per well) and the cells were incubated for an additional 6 hours protected from light. Afterward, the plates were centrifuged at 500 x g for 5 minutes, and the supernatant was discarded. The formazan crystals formed in the wells were dissolved with 200 µL of dimethyl sulfoxide (DMSO), and absorbance at 570 nm was measured using a SpectraMax M3 microplate reader (Molecular Devices). Cell viability was then calculated as 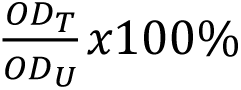 where OD_T_ represents optical density of the nanocarrier treated sample and OD_U_ corresponds to optical density of the untreated sample (n=3).

### Nanocarrier uptake studies

A total of 100,000 HUVECs or primary human Schlemm’s canal cells were plated in each well of 24-well polystyrene plates (Falcon) and allowed to adhere overnight at 37 °C with 5% CO_2_. The cells were exposed to nanocarriers for 2 hours, with all incubations conducted at 37 °C and 5% CO_2_. Following the nanocarrier treatment, cells were harvested by mechanical scraping and then stained with fixable Zombie Aqua viability dye (Biolegend, San Diego, CA) for 20 minutes at 4 °C to evaluate cytotoxicity by flow cytometry. Flow cytometry was performed using a BD LSRFortessa 6-Laser Flow Cytometer, recording 10,000 single-cell events per sample. Data were analyzed using FlowJo software. Median fluorescence intensity (MFI) was normalized by subtracting the average MFI from untreated cells, and this normalized value was used to assess nanocarrier uptake.

### Intracameral injection of 4OHT nanocarriers in mice

Mice were anesthetized with isoflurane and topical 0.5% proparacaine was provided for local analgesia. A 35G beveled needle (World Precision Instruments, NF35BL) was inserted into the anterior chamber and 3 μL nanocarrier solution was injected at a flow rate of 100 nL/min for 30 minutes using a micro digital syringe pump (UMP3, World Precision Instruments, Sarasota, FL). Following injection, topical ophthalmic antibiotic (Neo-Poly-Bac, Bausch + Lomb, Chapel Hill, NC) was applied to both eyes. Mice then received a single subcutaneous injection of Meloxicam (20mg/kg B.W.) for analgesia. A second administration of topical antibiotic was provided 24 hours later.

### Tonometric IOP measurements

Mice were anesthetized with ketamine/xylazine cocktail (ketamine 112.5 mg/kg, xylazine 2.5 mg/kg), and IOP measurements were taken 5–7 minutes post-anesthesia with a Tonolab rebound tonometer (iCare, Vantaa Finland). Individual Tonolab measurements represent averages of six individual recordings and IOP values reported in the manuscript are averaged from three measurements.

### In vivo visible light OCT imaging and Schlemm’s canal measurement

Visible light OCT (vis-OCT) uses shorter wavelengths than commonly used OCTs, thereby allowing for higher axial resolution and higher backscattering contrast in tissue. vis-OCT was performed using a custom-built OCT system mounted on a robotic arm as previously described to image the full 360° of the AHO pathway^64^. The system operated over a 510–610 nm spectral range, providing a theoretical axial resolution of 1.3 μm and a lateral resolution of 9.4 μm in tissue^64,65^. To achieve optimal sampling density while maintaining a compact imaging field, the lateral field of view for each volumetric scan was configured to 1.58 mm × 1.58 mm.

*In vivo* imaging was performed on mice under general anesthesia. Mice were anesthetized with an intraperitoneal injection (10 mL/kg body weight) of a ketamine/xylazine cocktail (11.45 mg/mL ketamine and 1.7 mg/mL xylazine in saline). To expose the limbal region for imaging, we made small relaxing incisions at the nasal and temporal canthi and inserted a circular eyelid speculum beneath the eyelids. During the imaging, a robotic arm (Meca500, Mecademic Inc.) precisely rotated the vis-OCT imaging head around the optical axis of the eye, enabling acquisition of eight volumetric vis-OCT datasets evenly distributed around the limbus at 45° intervals^65^. Each volume comprised 512 A-lines per B-scan and 512 B-scans per volume, collected using a temporal speckle-averaging acquisition protocol in which each B-scan was repeated twice per volume to improve signal-to-noise ratio and reduce speckle variation^66^. The imaging beam was delivered at an A-line rate of 75 kHz with an incident optical power of 1 mW at the cornea. All quadrants of the mouse eye were imaged at the same height relative to the imaging beam to ensure uniform sampling of the circumferential Schlemm’s Canal structures.

For quantitative assessment of Schlemm’s canal lumen dimensions, we analyzed two representative B-scans per vis-OCT volume, resulting in 16 B-scans per eye across the whole 360° imaging sequence. Each selected B-scan was chosen based on optimal visualization of the lumen and surrounding anatomical landmarks. The canal boundaries were manually segmented using MATLAB’s Volume Segmenter application, and we delineated the canal’s area using the paintbrush tool. The cross-sectional area of Schlemm’s canal was calculated for each segmented B-scan.

### Whole-mount imaging of mouse Schlemm’s canal

SC imaging was performed as previously described^67^. Briefly, whole globes were enucleated, fixed (2% formaldehyde in PBS, pH 7.5), and dissected to remove the lens and retina. Globes were then incubated overnight in lysis/blocking buffer (5% donkey serum, 3% bovine serum albumen in TBS pH 7.5 containing 0.5% triton X 100) prior to an additional overnight incubation in primary antibodies diluted in additional blocking buffer. Eyes were then washed, incubated with alexafluor-conjugated secondary antibodies and mounted for imaging. Images were captured using a Nikon A1R or Zeiss LSM 680 confocal microscope equipped with a 20x objective. 10-15-image Z stacks were obtained, and maximum intensity projections generated using Fiji software are shown in the manuscript. For quantification, 2-4 stacks were collected at intervals around the circumference of the eye and total fluorescence projections obtained using the “Sum Slices” function in ImageJ Fiji. Quantification of background subtracted protein expression and canal area were obtained from these images and averaged to obtain the values reported in the manuscript. Primary antibodies used: Goat anti human PROX1 (R&D systems AF2727), rabbit anti mouse PROX1 (Sigma-Aldrich-Chemicon, AB5475), goat anti mouse FLT4 (R&D Systems AF349), rat anti mouse PECAM1 (BD Biosciences AB397095), goat anti human PECAM1 (R&D Systems AF3628).

### Light and electron microscopy of mouse Schlemm’s canal

Mice were anesthetized using ketamine/xylazine cocktail and blood was cleared via cardiac perfusion at a flow rate of 15 ml/minute (PBS containing 1 mg/ml lidocaine and 10U/ml heparin). Following clearing, 50 ml of modified Karnovsky’s fixative (2% paraformaldehyde and 2.5% glutaraldehyde in 0.1M phosphate buffer, pH 7.4) was infused at the same flow rate. Upon completion of cardiac perfusion, eyes were enucleated, a small incision made to allow fluid transfer and submerged in additional fixative for 24 hours. Radial wedges containing trabecular meshwork and Schlemm’s canal were then collected from the nasal, inferior, superior and temporal quadrants and prepared for ultramicrotomy and imaging as previously reported^68^.

Briefly, wedges were post-fixed (1% OsO₄, 1% La(NO_3_)_3_, 2h) before staining with 1.5% uranyl acetate. Tissues were then dehydrated, washed with propylene oxide and embedded in Epon-Araldite using standard procedures. Gross Schlemm’s canal morphology was analyzed in toluidine blue-stained 4 μm semithin sections by a masked investigator. Images were analyzed using ImageJ. Reported values represent averages of 7-8 sections per eye, with 1-2 sections collected from each quadrant. For electron microscopy, ultrathin sections (70-80 nm) were prepared and imaging was performed using a JEOL JEM-1400Flash transmission electron microscope equipped with a NanoSprint43 cMOS camera.

### Statistical analysis and figure preparation

Statistical analysis was performed using Prism 10.6 software (Graphpad Software LLC). Tests used to obtain p values reported in the manuscript are described in figure legends. For all figures * indicates p < 0.05, ** p < 0.01, *** p < 0.001. Figures were prepared using Prism 10.6, Illustrator 29.8.1 and InDesign 20.5 software (Adobe Inc). Image analysis and quantification was performed using ImageJ Fiji ^69^ and Qupath Software. Except where noted, all error bars shown in figures indicate SEM.

## DISCUSSION

### Mechanism of IOP modulation by Schlemm’s canal

It has been well-known since at least the 1950s^6^ (and much longer to some^7^), that the ocular hypertension characteristic of primary open-angle glaucoma is due to increased resistance to the flow of aqueous humor through the outflow pathway in the eye. While cause of this increased resistance is still incompletely understood, several factors implicate the inner wall region of Schlemm’s canal including a decreased pore density^9,10^, increased cellular^11^ and tissue stiffness^12,70^, and an increased pressure drop in this region^12^. As there is also evidence that Schlemm’s canal endothelium modulates outflow resistance and thereby IOP in the normal eye^8^, here, we focused on how this modulation might occur.

Schlemm’s canal originates through sprouting angiogenesis from blood-filled capillaries in the limbal and iridocorneal angle regions of the eye^21,26^. PROX1 expression begins concurrently with the process of canal lumenization, and initiates expression of FLT4 and other lymphatic markers. The molecular identity of true lymphatic vessels is maintained by a signaling loop of PROX1 and FLT4^48,56^. Without continuous PROX1 transcriptional activity, lymphatic endothelial cells revert towards a venous phenotype and eventually degenerate^56^, consistent with the loss of limbal lymphatic capillaries that we observed in *Prox1* knockout eyes. Conversely, while specific deletion of *Prox1* from Schlemm’s canal markedly increased IOP (Fig. 4b-c), we did not observe canal degeneration, even over timelines up to 6 months, and measurements by *in-vivo* visible-light OCT, *ex-vivo* whole-mount confocal imaging, and plastic cross sections did not reveal any reduction in canal size (Fig. 4d-i, Fig. 5).We further showed relevance of this model to the human eye by confirming the presence of PROX1 in human Schlemm’s canal (Fig. 1b), as predicted by single cell RNA sequencing^42,45^ and in agreement with the findings of Aspelund et al.^30^ but contrary to the report of Birke et al.^46^. Furthermore, we observed reduced FLT4 expression in primary human Schlemm’s canal endothelial cells and in mouse eyes following loss of PROX1, but no increase in IOP following endothelial-specific *Flt4* deletion. Together, these findings suggested PROX1 regulated FLT4 expression in the canal as it does in lymphatics and that the PROX1-mediated hybrid phenotype was essential for IOP homeostasis. However, the PROX1-FLT4 feedback loop is not required for survival of Schlemm’s canal endothelial cells. Instead, lacking PROX1, Schlemm’s canal could indefinitely revert to a less functional venous phenotype.

Lymphatic capillaries are less stiff^71^ and more permeable^22^ than veins, characteristics central to the function of Schlemm’s canal. How these characteristics are regulated by PROX1 in the canal remains unknown, but our *in-vitro* studies provided compelling clues. Following siRNA-mediated *PROX1* knockdown, expression of *PTPRB,* the gene encoding the TEK and VE-Cadherin-regulating phosphatase VE-PTP, was markedly increased (Fig. 1f). Deletion of *Ptprb* in mice or inhibition of VE-PTP function in mice, rabbits and humans has been shown to lower IOP^72–74^, and VE-PTP is a key regulator of endothelial eNOS activity, acting via the Tie2/AKT signaling pathway ^75^. Rare variants in *PTPRB* are also associated with reduced glaucoma risk^76^. *Ptprb*/VE-PTP is highly expressed in blood vascular endothelial cells but is absent in lymphatics^77^, consistent with transcriptional suppression by PROX1.

### Schlemm’s canal’s phenotypic plasticity highlights new approaches for glaucoma therapy

Reversion of the *Prox1* knockout Schlemm’s canal to a vein-like phenotype was mirrored by our observation that PROX1 expression was negatively correlated with age in human eyes (Supplemental Fig. 2a) and by a report that PROX1 expression was reduced by over 50% in mouse Schlemm’s canal by one year of age while blood vascular markers were upregulated^36^.

Following IOP, aging is the most important glaucoma risk factor^78^. However, why the aged eye is more vulnerable remains poorly understood. Combined with the finding of Gaasterland et al.^51^ that outflow resistance increases with age, these findings suggest that increased risk of ocular hypertension is associated with loss of PROX1 expression from Schlemm’s canal in aged eyes and reversion towards a venous phenotype. While we found that lymphatic identity was not required for Schlemm’s canal endothelial cell survival, it had a critical role in function. As PROX1 expression decreases with age, reversion away from the outflow-optimized lymphatic fate may be a cause of age-related increase in outflow resistance. If true, these findings suggest that re-programming the aged canal to restore its functional hybrid phenotype would be an effective long-term strategy for enhancing outflow and providing protection from the ocular hypertension characteristic of primary open-angle glaucoma.

### Schlemm’s canal-specific Prox1 knockout mice as a new model of mouse ocular hypertension

Rodent genetic models of ocular hypertension have provided invaluable insights into glaucoma pathogenesis, genetics and treatment. However, many existing models are developmental in nature, exhibit very high IOP^79^ or are associated with ocular inflammation that complicates interpretation ^80^. Other models, such as *MYOC ^Y437H^*transgenic mice, have been reported to lose pressure elevation over time, limiting their usefulness for long-term studies^81,82^. Here, we report that Schlemm’s canal specific *Prox1* knockout mice exhibit ocular hypertension within 4 weeks of induction. IOP was increased by ∼5 mmHg in treated eyes, as compared with contralateral control eyes injected with empty nanocarriers, and elevated IOP was maintained for at least 6 months. As in human ocular hypertension, we did not observe gross structural abnormalities in angle morphology. Moreover, as PROX1 expression was negatively correlated with age in mouse^36,47^ and human eyes our data suggest that Schlemm’s canal specific *Prox1* knockouts are a highly relevant model of ocular hypertension that recapitulate aspects of canal aging for future studies examining the impact of elevated IOP on retinal function or testing IOP-lowering drugs.

### 4OHT nanocarriers are a valuable new tool for targeted gene deletion

By facilitating tissue and time-dependent gene recombination, cre-expressing mouse strains have revolutionized animal research. However, these tools depend on existence of promotors with appropriate tissue selectivity and other limitations, such as cellular toxicity of Cre recombinase expression and variable or inconsistent patterns of deletion, are increasingly appreciated. This underscores the need for new strategies with higher tissue selectivity and fewer confounding phenotypes^40,83^.

Here, we demonstrated that combining the well-validated endothelial *Cdh5*-CreERT2 mouse strain with tissue specific 4OHT delivery via a targeted non-inflammatory nanocarrier ^84–87^ allowed robust gene deletion and increased tissue specificity without ocular toxicity. While adeno-associated viruses (AAV) have been used in similar applications to generate models of eye disease^35^ and for gene therapy^88^, ocular inflammation has been observed in patients^89–91^ and rodent models^92^ following AAV administration. In contrast, the PEG-*b*-PPS platform is a non-inflammatory delivery system^93^ that can be customized for cell-specific targeting^84–87^ and adapted to deliver a wide array of cargos and combinations thereof^84,94,95^.

In addition to their small diameter that facilitated passage into the canal (∼22 nm), PEG-*b*-PPS nanocarriers are highly efficient at intracellular delivery^85,96^, improving therapeutic effects at lower concentrations for encapsulated payloads. This approach enhanced 4OHT efficacy for driving recombination in nearly all targeted Schlemm’s canal cells, allowing us to observe *Prox1*-dependent pressure elevation while maintaining monodisperse particle size and avoiding systemic effects of pan-endothelial *Prox1* deletion. The 4OHT-loaded PEG-*b*-PPS nanocarriers hold promise as a tool for achieving customizable and geographically targeted investigation of gene expression in other tissues and models, with observed responses being attributed to specific gene manipulation instead of non-specific inflammation or toxicity from tamoxifen or the delivery vehicle itself.

### Summary

Schlemm’s canal-specific *Prox1* deletion in adult mice leads to long-lasting ocular hypertension without canal degeneration or morphological changes. These findings demonstrate the importance of the canal’s lymphatic characteristics in the maintenance of normal aqueous humor outflow resistance and highlight the phenotypic plasticity of Schlemm’s canal as a target for next-generation therapies for ocular hypertension. In addition to these mechanistic insights, our studies also demonstrate the power of Schlemm’s canal-specific 4OH-tamoxifen nanocarriers for inducing robust, canal specific gene deletion without systemic phenotypes, as well as introduce canal specific *Prox1* knockout mice as a new model of primary open angle glaucoma.

## ACKNOWLEDGEMENTS

This work was supported by NIH R01 R01EY033813 (To MJ and EAS), a Northwestern University Catalyst Grant (To MJ and BRT), and the Christina Enroth-Cugell Graduate Research Award (To SLO). BRT also received support from NIH R01 R01EY032609 and Brightfocus Foundation grant M2021018N. Confocal imaging was performed at the Center for Advanced Microscopy of the Feinberg School of Medicine, supported by NCI CCSG P30 CA060553. We also acknowledge support from George M. O’Brien kidney core grant P30 DK114857, and a challenge grant from Research to Prevent Blindness to the Feinberg School of Medicine Department of Ophthalmology.

Authors are grateful to Drs. Guillermo Oliver and Susan E. Quaggin (Northwestern University Feinberg School of Medicine) for the generous gifts of *Prox1*-floxed and *Flt4*-floxed mice, respectively, and for many valuable discussions. We are also indebted to Dr. W. Daniel Stamer (Duke University School of Medicine) for the gift of primary human Schlemm’s canal endothelial cells. Technical support was provided by Sol Misener and Kyron McAllister, without whom this study would not have been possible.

## AUTHOR CONTRIBUTIONS

BRT, MJ and EAS conceived of the study. SLO, BRT, MJ, EAS, HG, HFZ, ZY and HLL designed experiments and analyzed results. SLO, MPV, HLL, NCM, PL, ZY, HJL and BRT conducted experiments. SLO, BRT, MJ and EAS wrote the manuscript. All authors contributed to editing the manuscript and validation of results.

---

**Supplemental Figure 1.**
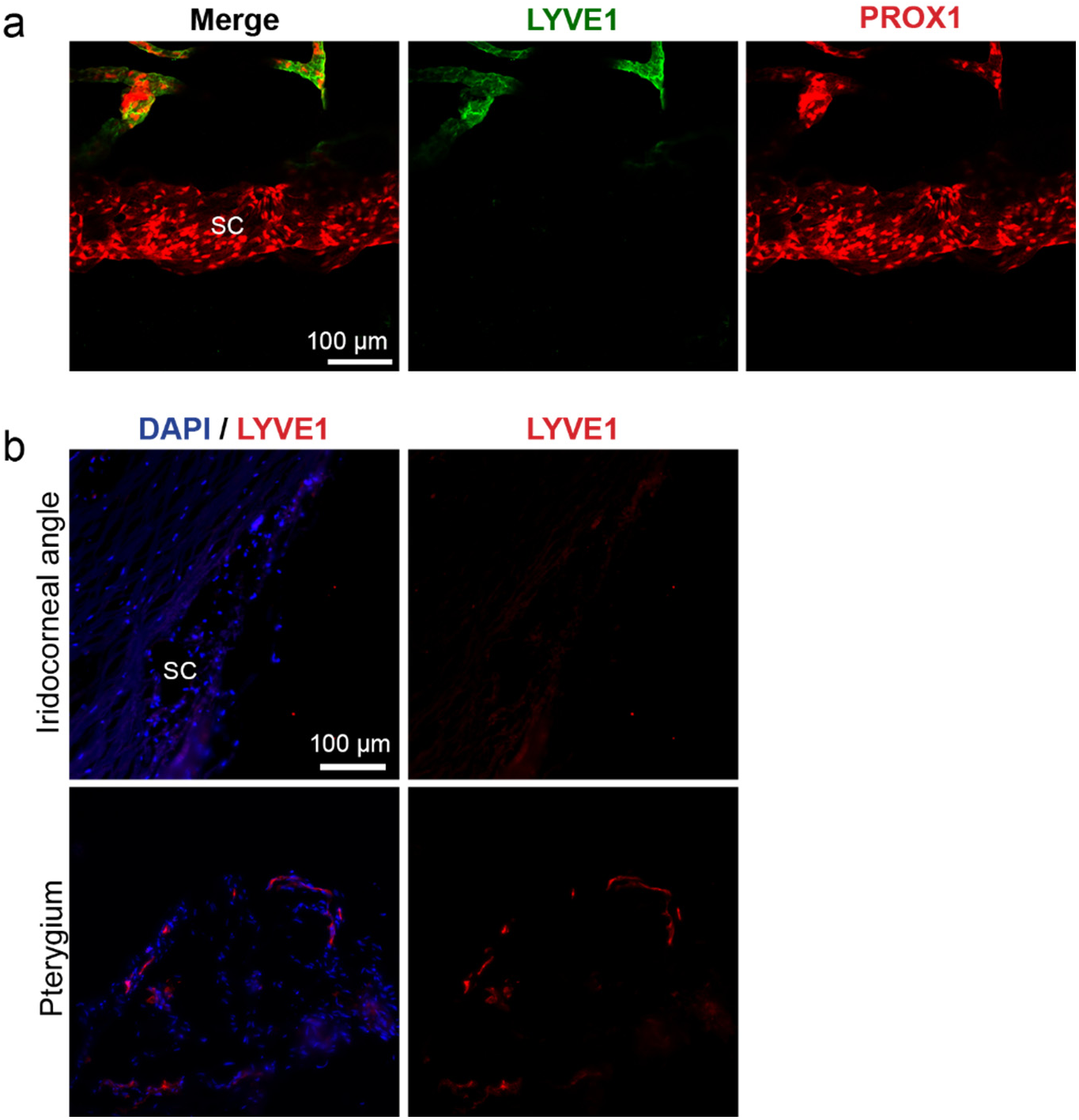
The lymphatic protein LYVE1 is not expressed in mouse or human Schlemm’s canal (SC). (**a**) Whole mount immunostaining of mouse S, and (**b**) cryopreserved cross-sections of human SC revealed no expression of the common lymphatic marker LYVE1, although robust expression was observed in adjacent lymphatic vessels of the mouse eye and in human pterygium tissue.

**Supplemental Figure 2.**
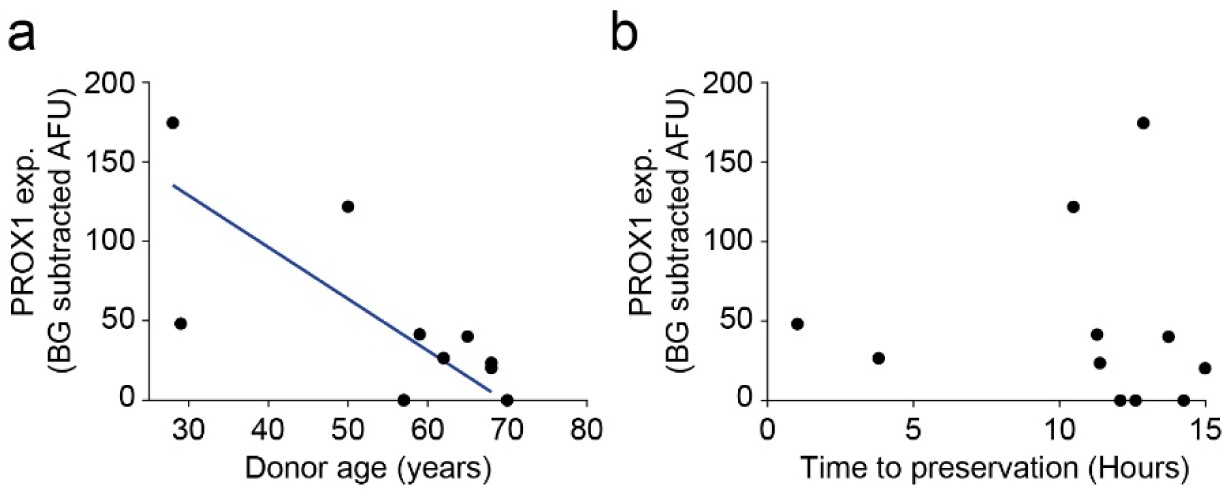
PROX1 expression was negatively correlated with age in human corneal rim cryosections. Results from multivariate regression of nuclear PROX1 immunostaining intensity against donor age and time to preservation (Full model, p=0.028, adjusted R^2^ = 0.4887, n = 10 donors) (**a**) PROX1 staining intensity within ERG-positive Schlemm’s canal endothelial nuclei was negatively correlated with age (age, p<0.01) in human corneal rim sections. (**b**) PROX1 staining intensity did not decrease with time from death to tissue preservation for eyes preserved in less than 15 hours of death (time to preservation, p>0.6).

**Supplemental Figure 3.**
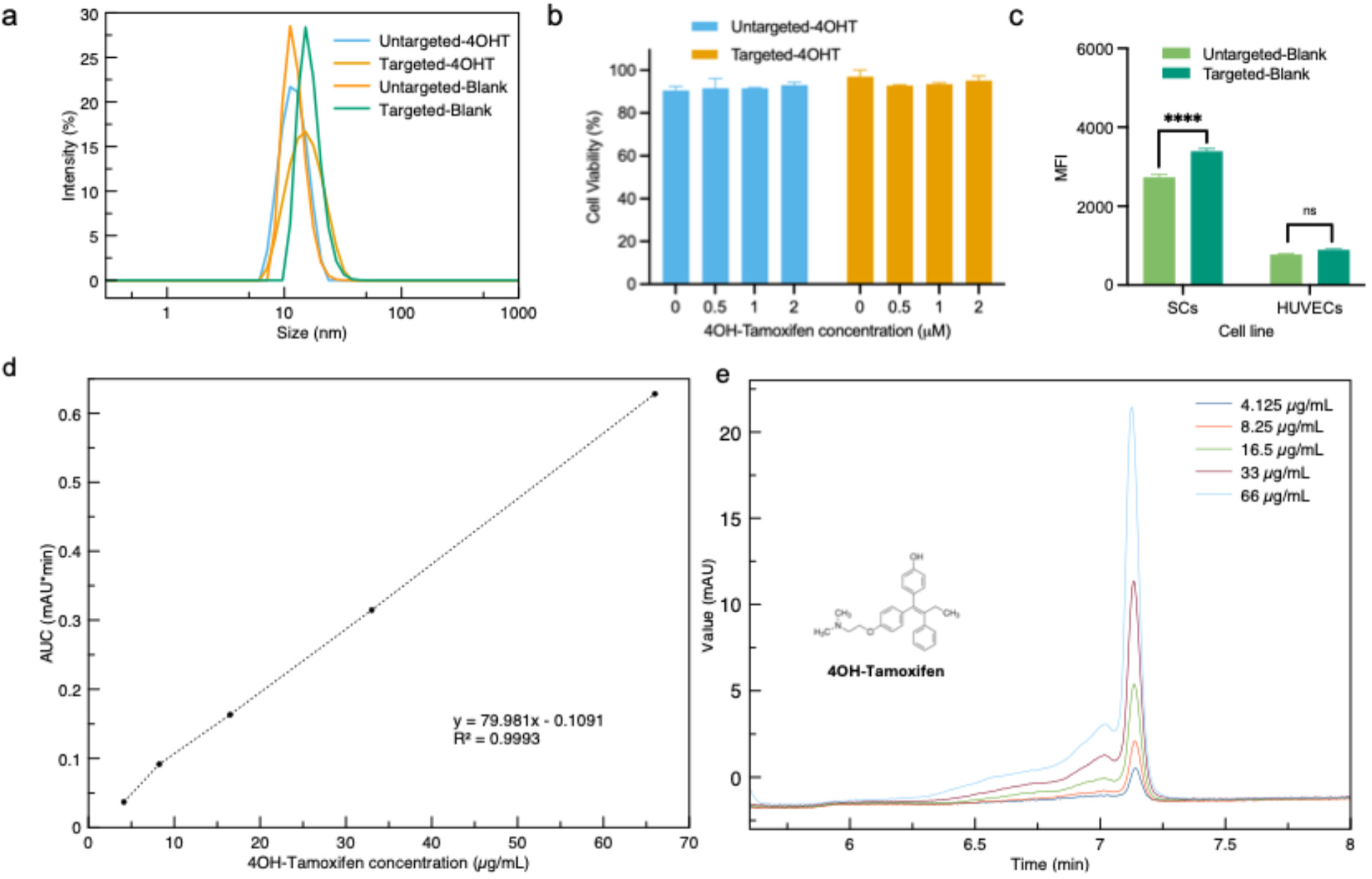
(a) Size distribution of 4 different targeted or untargeted samples containing 4OH-Tamoxifen fabricated with cosolvent evaporation. (**b**) MTT assay revealed that HUVEC cell viability was not affected by a 4h incubation with either 4OH-Tamoxifen targeted or untargeted nanocarriers. (**c**) Decoration with a VEGFC-derived targeting peptide increased uptake of DiI-labeled blank nanocarriers by primary human Schlemm’s canal endothelial cells but not HUVECs. The mean fluorescence intensity (MFI) *±* s.e.m. (n=4) is displayed. Significance was determined by ANOVA followed by Tukey’s multiple comparisons test, 5% significance level was used for all statistical tests. ****p<0.0001. (**d**) Standard curve of serially diluted free form 4OHT. A C18 XDB-Eclipse column (Agilent) was used with a static mobile phase of acetonitrile and 0.1% (v/v%) TFA water (85:15). (**e**) HPLC chromatograms of 4OH-Tamoxifen that referenced a reproducible 4OHT concentration series for standard curve calibration of High-Performance Liquid Chromatography (HPLC) measurements. A linear regression model fitted to the data: y = 79.981x + 0.1091, r^2^ = 0.9993 to determine the concentration of 4OH-Tamoxifen of the samples.

**Supplemental figure 4.**
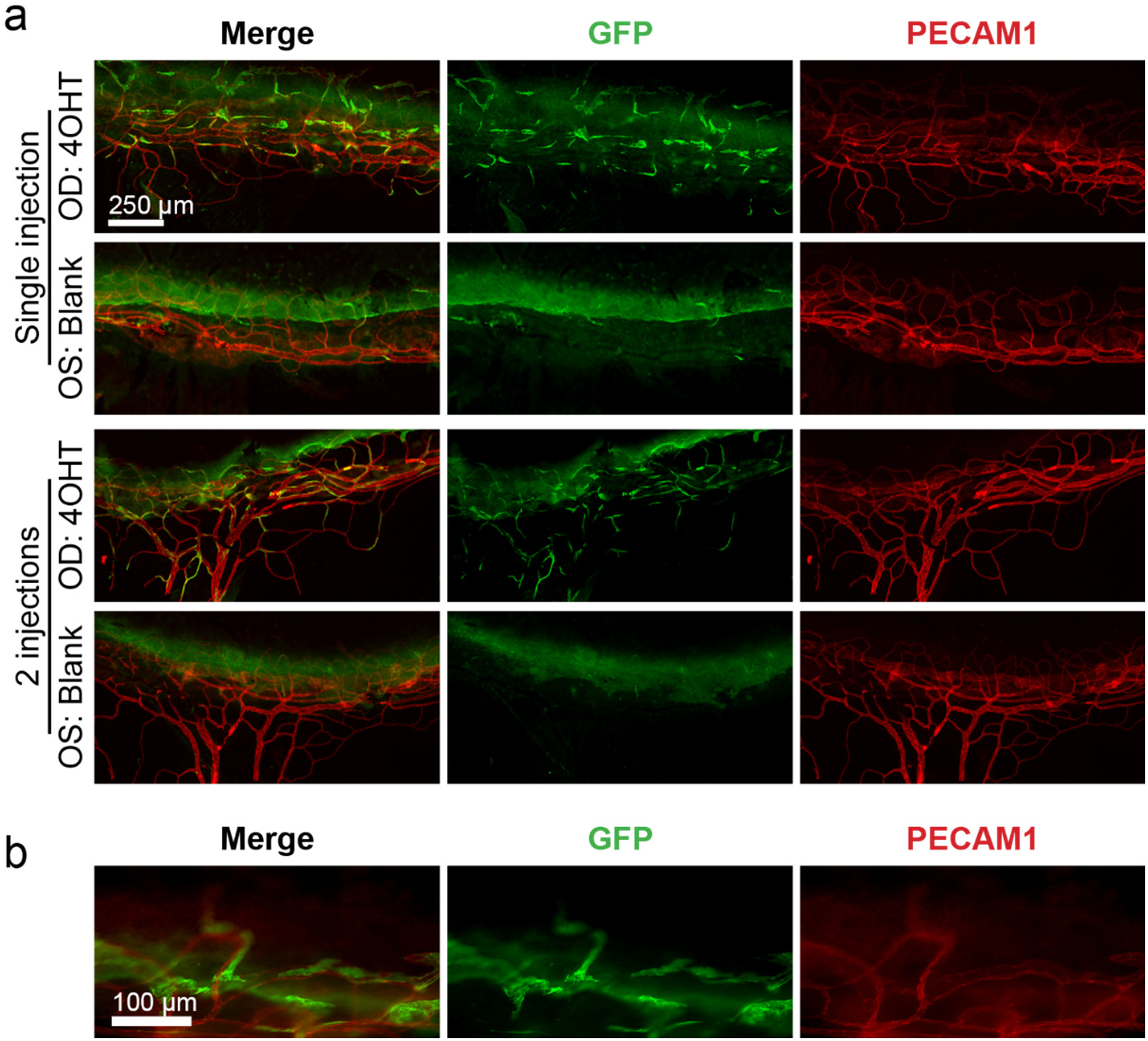
Sporadic recombination was observed in distal outflow and limbal blood vessels of eyes treated with 4-OH-tamoxifen loaded, Schlemm’s canal targeted nanocarriers. (**a**) 7 days after intracameral nanocarrier delivery, low levels of sporadic cre-mediated recombination (determined by GFP fluorescence) was observed in *Rosa26*^mTmG^; *Cdh5*-CreERT2 mice treated with 4OHT nanocarriers. Similar levels of recombination were seen in eyes receiving one or two nanocarrier infusions, while very few recombined cells were observed in contralateral control eyes that received matching injections of empty control nanocarriers. (**b**) Recombination was also observed in endothelial cells of limbal lymphatic capillaries of some eyes receiving targeted 4OHT nanocarriers.

**Supplemental figure 5.**
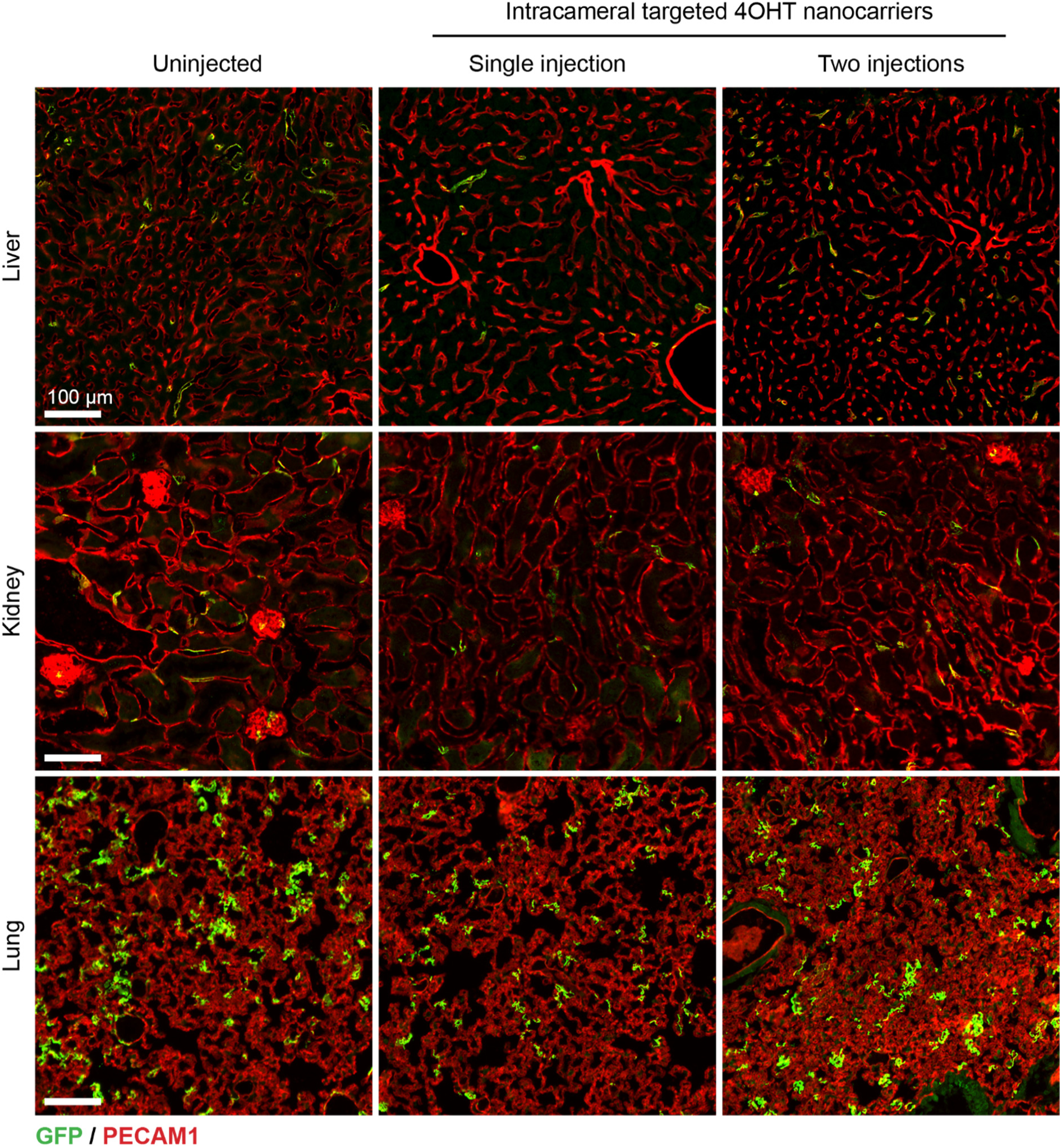
Outside of the eye, no increase in cre-mediated recombination was observed following intracameral injection with 4OHT loaded nanocarriers. Cryosections were prepared from liver, kidney and lung tissues of *Rosa26*^mTmG^; *Cdh5*-CreERT2 mice seven days after intracameral injection with 4OHT-loaded Schlemm’s canal targeting nanocarriers, and cre mediated recombination was analyzed by GFP fluorescence. While some recombination was observed in endothelial cells of all organs, number of recombined endothelial cells was similar in untreated and nanocarrier-treated mice, suggesting this was due to cre leakiness and not nanocarrier activity.

**Supplemental figure 6.**
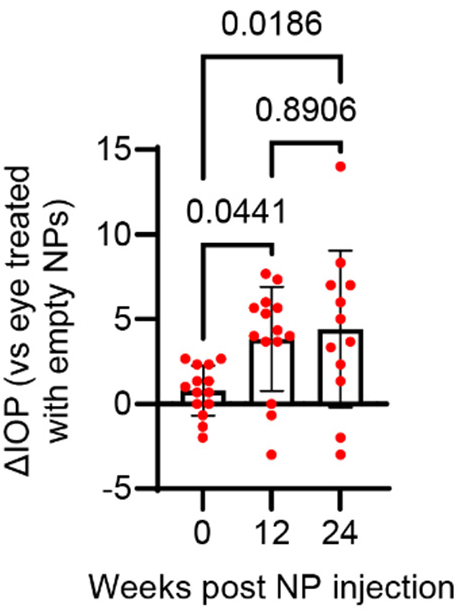
Prolonged IOP elevation in a second cohort of *Prox1*^flox/flox^; *Cdh5*-CreERT2 mice after 4-OH tamoxifen nanocarrier treatment. IOP elevation was maintained in Cre-positive mice until animals were euthanized 24 weeks after nanocarrier-mediated PROX1 deletion. Error bars indicate ±SEM, reported p values were obtained using a 1-way ANOVA followed by Tukey’s multiple comparison test. n = 14 mice.

**Supplemental figure 7.**
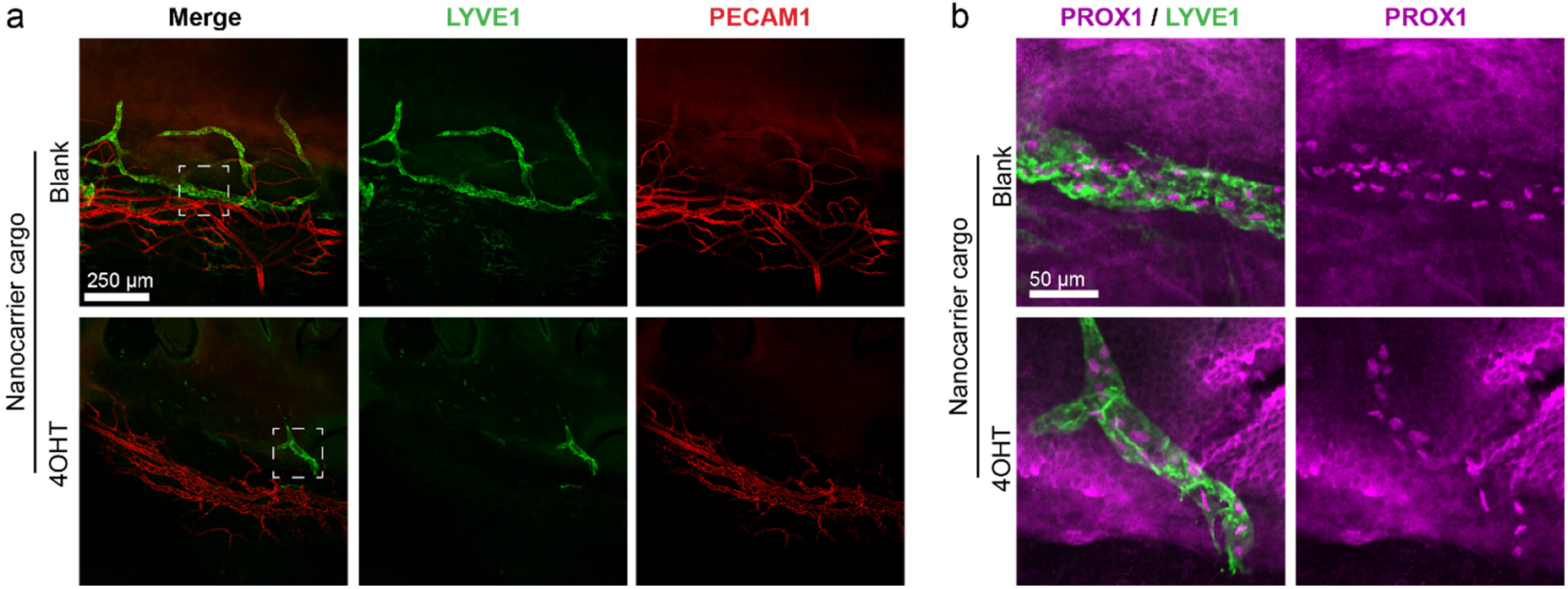
Degeneration of limbal lymphatic vessels was observed in *Prox1*^flox/flox^; *Cdh5*-CreERT2 eyes following 4OH-tamoxifen nanocarrier induction. (**a**) Compared with the organized pattern of limbal lymphatic capillaries in *Prox1*^flox/flox^; *Cdh5*-CreERT2 eyes treated with blank control nanocarriers, confocal microscopy at the level of the limbal vascular network revealed degeneration of limbal lymphatics in contralateral eyes receiving 4OHT nanocarriers. (**b**) Normal PROX1 expression was observed in remaining lymphatic endothelial cells of 4OHT-treated eyes. Dashed white square in (**a**) indicates regions of detail highlighted in (**b**).

